# Quantifying the regulatory role of individual transcription factors in *Escherichia coli*

**DOI:** 10.1101/2021.01.04.425191

**Authors:** Sunil Guharajan, Shivani Chhabra, Vinuselvi Parisutham, Robert C. Brewster

**Affiliations:** Program in Systems Biology, University of Massachusetts Medical School, Worcester, MA 01605; Department of Pharmacological Sciences, Icahn School of Medicine at Mount Sinai, New York, NY 10029; Department of Microbiology and Physiological Systems, University of Massachusetts Medical School, Worcester, MA 01605

**Keywords:** Systems biology, Synthetic biology, Transcription regulation

## Abstract

Gene regulation often results from the action of multiple transcription factors (TFs) acting at a promoter, with a net regulation that depends on both the direct interactions of TFs with RNA polymerase (RNAP) and the indirect interactions with each other. Here we measure the fundamental regulatory interactions of TFs in *E. coli* by designing synthetic target genes that isolate the individual TFs regulatory effect. Using a thermodynamic model, the direct regulatory impact of the TF on RNAP is decoupled from TF occupancy and interpreted as acting through two mechanisms: (de)stabilization of RNAP and (de)acceleration of transcription initiation. We find the contributions of each mechanism depends on TF identity and binding location; for the set of TFs profiled, regulation immediately downstream of the promoter is insensitive to TF identity, yet these same TFs regulate by distinct mechanisms upstream of the promoter. Strikingly, we observe two fundamental regulatory paradigms with these two mechanisms acting coherently, to reinforce the observed regulatory role (activation or repression), or incoherently, where the TF regulates two distinct steps with opposing effect. This insight provides critical information on the scope of TF-RNAP regulation allowing for a stronger approach to characterize the endogenous regulatory function of TFs.

## Introduction

Transcriptional regulation of gene expression is one of the major mechanisms by which cells respond to cues and stimuli. Transcription factors (TFs) perform this regulation through binding to the DNA around the promoter to alter the rate of transcription from individual genes [1, 2]. The regulatory DNA of each gene is distinct and can involve several to dozens of TF binding sites arranged in specific architectures to achieve the desired expression level. However, predicting the level of gene expression based on the regulatory architecture of a gene remains a central challenge in the field [3, 4, 5, 6, 7].

The genomics era has enabled multiple techniques capable of determining where a TF will bind and with what specificity [8, 9, 10, 11]. Although this information is crucial for building occupancy-based models of gene regulation, there is still another critical component which is missing; the quantitative regulatory role of a TF, once bound, is often unclear. Historically, measurements of TF function are the result of knocking out the endogenous TF and observing how gene expression changes as a result. This serves its purpose in predicting the specific role of that TF on a given gene but offers less predictive power when examining regulation of other genes by that same TF or to different binding sites. These measurements of gene regulation are often entangled in indirect regulatory effects such as TF-TF interactions [12, 13], feedback [14, 15], physiological (*i*.*e*. growth rate) [16, 17] and off-target competitive effects of decoy binding sites or other genes in the network [18, 19]. Due to this, a single TF that binds at the same relative location on two different natural promoters can appear to have opposite regulatory roles. The entanglement between indirect and direct regulation likely contributes to this ambiguity and prevents the field from developing a basic intuition of TF regulatory function. The select few TFs that have arisen as “model TFs” such as LacI [20, 21, 22, 23, 24], AraC [25, 26, 27, 28], lambda repressor [29, 30, 31] and TetR [32, 33, 34, 35] have well studied regulatory function. Indeed, these TFs have been utilized for design of synthetic circuits with engineered purpose such as the creation of logic gates [34, 36], bistable switches [35, 37], oscillatory networks [38, 39, 5], synthetic enhancers [40, 41] and a host of other dynamic outcomes. Further characterizing the regulatory function of TFs beyond this small subset should provide a more complete toolset for broader synthetic design purposes.

Here we study the isolated regulatory function of a set of *E. coli* TFs in a system designed to remove the typical confounding factors of natural genes and quantify the direct regulatory effect of a TF based on factors such as TF concentration, binding affinity and binding location. Using a collection of strains where the average copy number of most TFs in the cell can be controlled, we measure the level of regulation of an individual TF acting on a synthetic promoter sequence. This promoter is designed to be regulated only by that TF, and it is targeted to a binding site whose location and sequence we control. To interpret this data, we use a thermodynamic model of gene regulation to parameterize TF regulatory function. In principle, the TF could exert its regulatory effect at any one of the distinct kinetic steps of the transcriptional process [42, 43, 44, 45] or on several of them simultaneously, and our model course-grains TF activity into two distinct modes of regulation. The first regulatory mode “stabilization” corresponds to stabilization (or destabilization) of the polymerase at the promoter by the TF. The second mode “acceleration” corresponds to a TFs ability to accelerate (or decelerate) the initiation of transcription when both the TF and polymerase are bound to the promoter. Using this model we infer the quantitative contribution from each of these modes in the data. Importantly, this process allows for the decoupling of properties that are extrinsic to the TF such as affinity to the operator binding site, the overall concentration of the TF or feedback in the network from the core regulatory role of the TF in modulating the steps of the transcription process.

We expect the regulatory parameters of a TF to vary based on the identity of the TF and the binding site location on the regulated promoter. In this study, we investigate the role of TF identity by measuring regulation of 6 TFs (AcrR, AgaR, ArsR, AscG, BetI, and CpxR) at two common binding locations: directly downstream of the promoter, where repression is commonly observed, and 61 bases upstream of the promoter, a site commonly associated with activation (although databases of regulatory interactions record roughly as many TFs repress at −61 as activate). We find that, despite the diverse nature of the TFs tested (five of the TFs are annotated repressors and one of them, CpxR, is a known activator), the regulation for all TFs immediately downstream is consistent with a form of repression that is set by the degree of occupancy of the TF at the promoter - independent of TF identity. This commonality across the TFs disappears when we measure the effect at -61, where the TFs exhibit different degrees of stabilization with both CpxR and AgaR engaging in significant stabilization of RNAP. To compliment this, we took CpxR and systematically quantified the contribution of the regulatory modes as a function of TF binding location and find that CpxR sets the degree of activation by engaging in two distinct regulatory paradigms. Binding locations that see strong activation have CpxR engaging in “coherent” regulation: the activation is enforced by stabilization and acceleration of RNAP. Locations with weak activation, however, have CpxR regulating the two modes oppositely by stabilizing RNAP yet slowing the rate of promoter escape - demonstrating that such “incoherent” regulation plays a useful role in allowing a single TF to generate a spectrum of regulatory responses emerging from the relative effects of the TF on these distinct steps.

## Results

### Thermodynamic model for single TF regulation

In order to deconvolve the role of TF copy number, binding affinity and binding location from the intrinsic regulatory interactions of the TF with polymerase we use a thermodynamic model of gene expression [46, 47, 48, 49, 50, 51, 52] where we consider only a single TF acting on an otherwise unregulated gene. Figure 1A shows the various promoter states considered in the model (left column) along with the relative probability of each state occurring (middle column) and the rate of expression from each promoter state (right column); the promoter can be unbound by TF and RNAP, bound by polymerase only, bound by TF only, or bound by both. The probability with which these states occur is a function of each molecules (polymerase and TF) binding affinity with their specific DNA binding sites (Δ_*ϵ*P_ and Δ_*ϵ* TF_) and the available number of each molecule in the cell (*N*_*p*_ and *N*_TF_). For the cobound state, we consider two distinct mechanistic influences of the TF on gene expression. The first effect represents altered stability of the polymerase at the promoter when TF is bound due to a favorable (activating) or disfavorable (repressive) interaction between TF and polymerase. As a result, the cobound state occurs with increased relative probability to the single bound state by a factor *β* (implying an energetic interaction of log(*β*) in units of *k*_B_*T*). The second parameter *α* represents the change in transcription rate when the TF and polymerase are cobound and is written as a multiplicative factor to the base expression rate of polymerase bound in the absence of TF; for example, *α* = 2 would imply that transcription initiation rate is doubled when the TF and polymerase are co-bound. In both cases the parameters represent increases to gene expression when greater than unity and decreases of gene expression when less than unity. Importantly, the parameters are not constrained and can, in principle, have opposing or compounding effects; *i*.*e*. this model allows for a TF that both stabilizes polymerase binding but slows the rate of transcription from that state resulting in apparent activation or repression depending on the relative strengths of those effects. The final parameter, *N*_NS_ is equated to the size of the genome in base pairs (4.6 ×10^6^) and is not varied in our experiments, for more details see [53, 48]. Further-more, the parameters related to polymerase binding can be simplified into a single parameters in our model as *P* = *N*_*p*_exp (-Δ*ϵ*_P_)*/N*_NS_.

**Figure 1:**
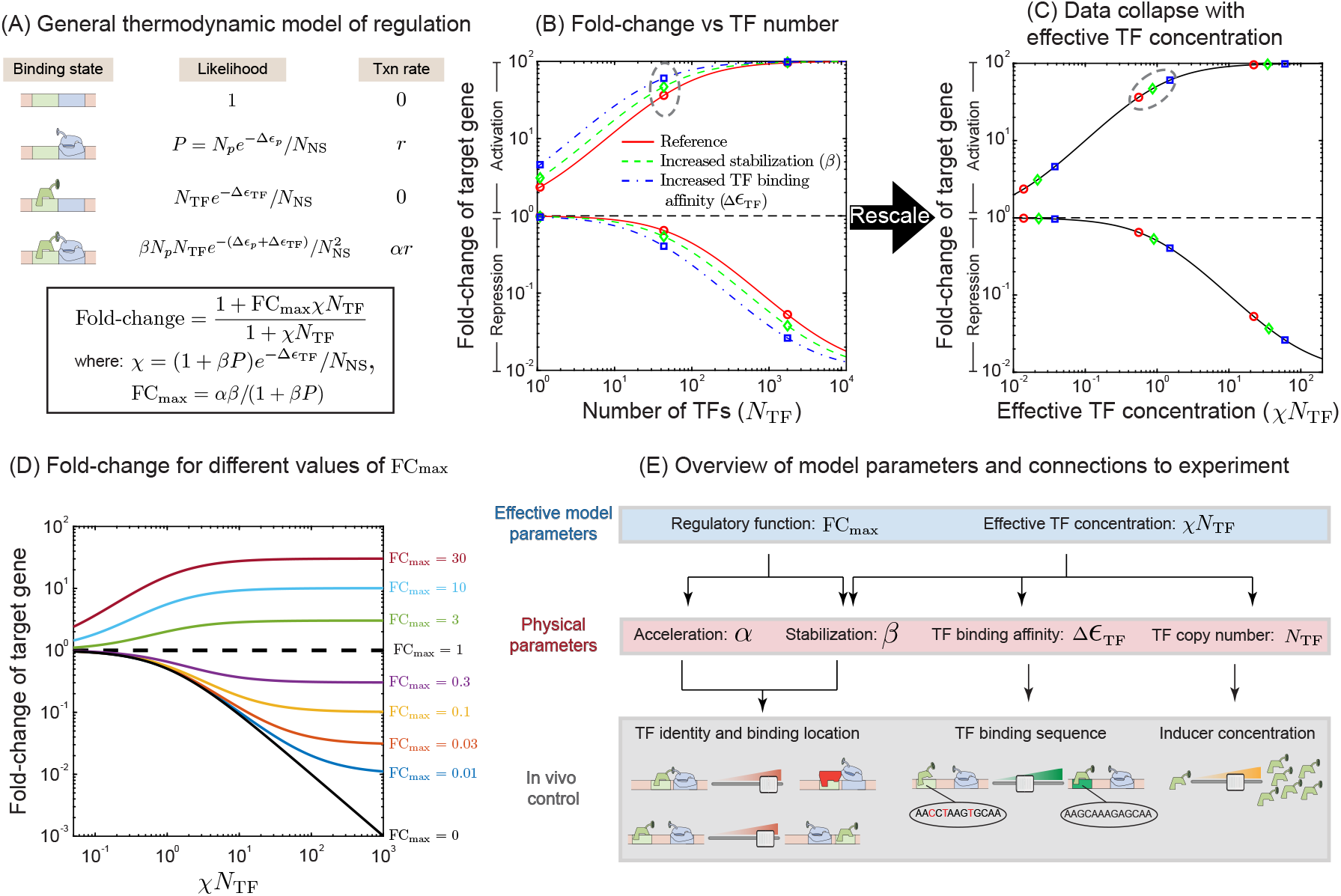
Thermodynamic modeling for measuring TF regulatory features. (A) The thermodynamic model for the target gene allows for four states: unbound by TF or RNAP, bound by RNAP, bound by TF or bound by both. The probability of each of these states occurring is listed in the middle column. The rightmost column shows the transcription rates in these states. (B) Fold-change vs TF copy number (*N*_TF_) for gene regulated by an activator (top set of curves) or a repressor (bottom set of curves). The blue and green curves have the same FC_max_ as the red curve but with increased stability (*β*, green curve) or TF binding affinity (Δ*ϵ*_TF_, blue curve). (C) Replotting the curves in (B) as a function of effective TF concentration (*χN*_TF_) demonstrates that each of the curves now falls onto a single “collapsed” curve defined by the effective parameters FC_max_ and *χN*_TF_. (D) Data collapse curve for a range of different FC_max_ values. (E) The relationship between theoretical parameters of the model and *in vivo* molecular details of the regulation.

The final expression, boxed below in Figure 1A (and derived in the Methods section), predicts the fold-change in gene expression of a target gene. Fold-change is defined as the expression level of the target gene in the presence of a number of TFs (*N*_TF_) divided by the expression level in the absence of that TF (*i*.*e N*_TF_ = 0). Fold-change greater than 1 signifies activation, while fold-change below 1 signifies repression. The fold-change equation is simplified by collecting the regulatory parameters into two effective parameters FC_max_ and *χ*. FC_max_ represents the fold-change when the number of TFs in the system are saturating; in the case of a repressor it is the minimum fold-change achievable and in the case of an activator it is the maximum fold-change achievable. Importantly FC_max_ depends only on a TFs degree of acceleration (*α*) and stabilization (*β*) and not TF binding affinity or concentration. The second term, *χ*, represents the rate at which the fold-change approaches FC_max_. This rate of approach depends on the TF binding affinity (Δ_*ϵ* TF_) and the degree to which the TF recruits/stabilizes RNA polymerase (1 + *βP*). These factors together with the number of TFs (*N*_TF_) can be thought of as an effective TF concentration.

These effective parameters are useful because they transform this system with many variables (TF and polymerase binding affinity, TF and polymerase number, degree of acceleration, degree of stabilization, *etc*) capable of producing a diverse range of response curves in the fold-change vs TF copy number space into a very simple system dictated by two fundamental quantities: the maximum fold-change FC_max_ and the effective TF concentration *χN*_TF_. This is demonstrated in Figure 1B where we plot fold-change against TF number for an activator (upper curves) and a repressor (lower curves) with FC_max_ = 100 and FC_max_ = 1*/*100 respectively. In each scenario we plot 3 colored curves, the red curve has no contribution from stabilization (*β* = 1). The blue curve is identical to the red curve except with slightly stronger TF binding affinity and the green is once again identical to the red curve except with significant contribution from stabilization (*β* = 10 with *α* adjusted to keep FC_max_ unchanged). Figure 1C demonstrates the fold-change as a function of effective TF concentration, *χN*_TF_. When plotted this way the data from all three curves collapses to a single curve that is determined entirely by the value of FC_max_, independent of specific values of *α* and *β*. Points with identical TF concentrations in Figure 1B now scatter on the collapsed curves and the green and blue points (which had higher values of *χ*) are further along the curve; a large value of *χ* hastens the approach to FC_max_ for the same TF concentration (*N*_TF_). Specifically, two TFs with similar net regulatory effect (similar FC_max_) that operate through different regulatory mechanisms, for instance one through strong acceleration (large *α, β* ≈ 1) and one through strong stabilization (large *β, α* ≈ 1), will trace out exactly the same curve in this space, however the strong stabilizer will have a higher effective TF concentration and thus move further along the curve given the same TF concentrations and binding affinities. In figure 1D, data collapse curves for a range of FC_max_ values is shown.

Figure 1E provides a hierarchical overview of model parameters and their relationship to controllable biological features of the *in vivo* system. The upper level shows the effective parameters FC_max_ and *χN*_TF_ which define the contour of regulatory curves like those in Figure 1D. These effective parameters are composed of a combination of the “physical” parameters on the second level of the diagram. These parameters correspond to basic features of the system such as numbers of molecules, affinities or interactions between molecules. The third level of the diagram shows the biological controls we have available to control the corresponding physical parameters. The approach we take below will be to profile the regulatory function and characterize the inherent regulatory parameters (*α* and *β*) of six TFs (AgaR, ArsR, AcrR, AscG, BetI, and CpxR) by controlling the copy number, binding location and binding sequence of each. A potential challenge in determining *α* and *β* stems from their connectedness in the effective parameters *χ* and FC_max_. Due to the fact that *χ* is proportional to (1 + *βP*), if the stabilization parameter *β* is much smaller than 1*/P*≈ 15 (measured previously for the promoter sequence used in our experiments [54]) then *χ* will no longer strongly depend on *β* because (1 + *βP*) ≈1. In cases such as this, which we label “weak stabilization”, we are left with only one effective parameter FC_max_ to determine the two regulatory parameters *α* and *β* and it is not possible to distinguish between regulation driven by a change in the transcription rate (acceleration/deceleration) or by modulation of polymerase occupancy at the promoter (stabilization/destabilization).

### Experimental measurements of individual TF regulatory function

To measure regulation by an individual TF in *E. coli* as a function of TF identity and binding position on the promoter we utilize synthetic techniques to create simple and controllable gene circuits. In this approach we have created *E. coli* strains, schematically illustrated in Figure 2A-B, where the endogenous copy of each studied TF is knocked out and reintroduced as a TF-mCherry fusion integrated into the genome at the *ybcN* locus (Figure 2A). Expression of the synthetic TF-mCherry promoter can be induced with anhydrotetracycline (aTc) (for details, see *Methods*). This system enables precise control of TF copy number (Figure 2D) which is measured either by wide-field fluorescent microscopy or flow cytometry (Figure 2C, see *Methods*). Using this system we are able to induce these TFs from leaky levels (several per cell) up to several thousand per cell; the full induction curve of each strain is shown in Figure 2D. Importantly, although we control the concentration of the TF, many TFs are capable of existing in distinct binding conformations that may alter the active TF number, the regulatory parameters (*α* and *β*), or both. For all but one of the TFs studied here, we expect that the TFs will always be active in our growth conditions, however BetI is inactivated by choline (which we do not control for) and as such we expect to have some fraction of BetI inactive at low TF concentrations [55, 56].

**Figure 2:**
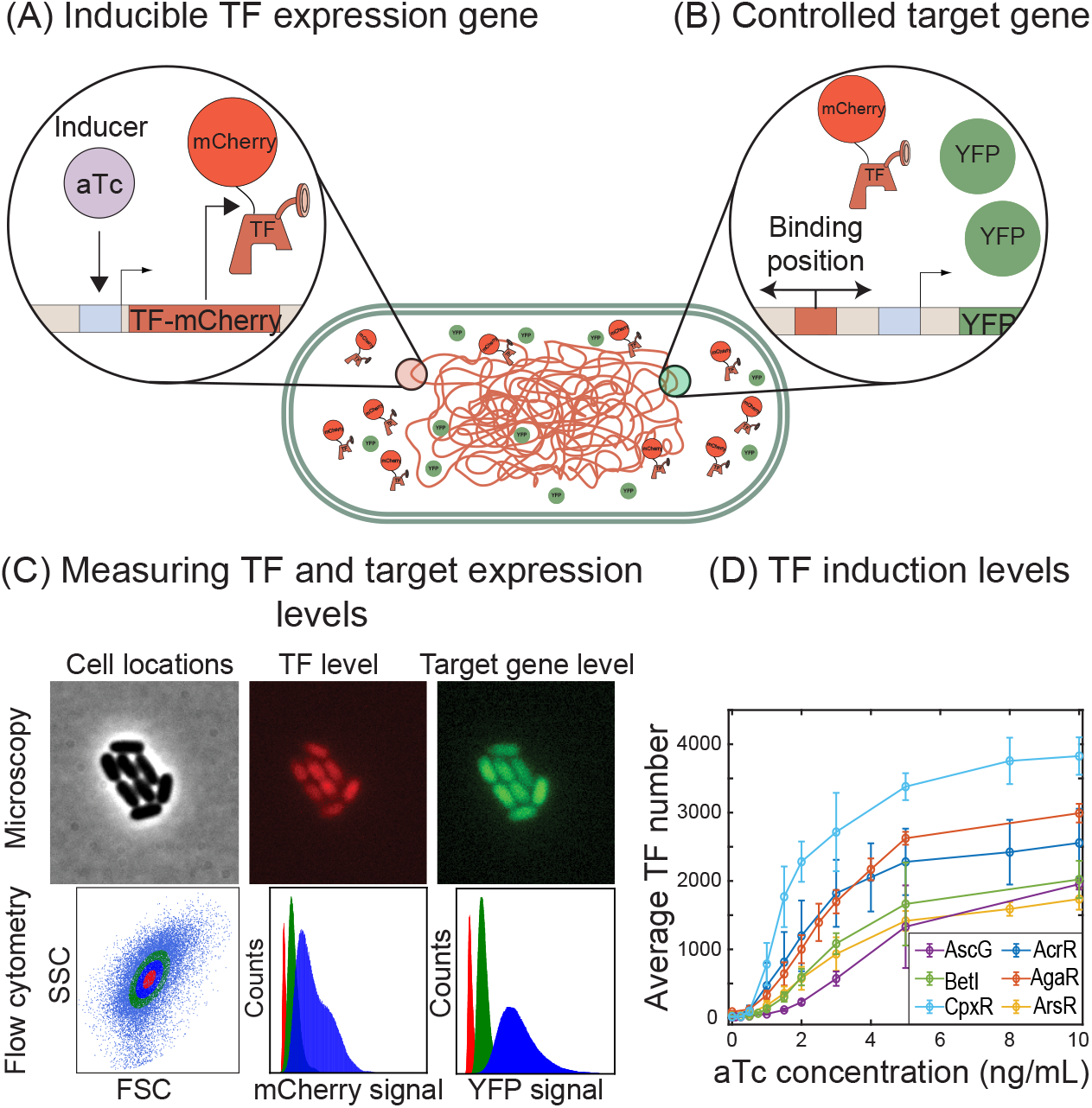
Experimental setup for measuring the TF position dependent regulatory profile. (A) The inducible TF expression strains consist of a set of base strains where the endogenous copy of any one of our 6 TFs knocked out and reintroduced as a TF-mCherry fusion at the *ybcN* locus expressed from an inducible *tet* promoter. (B) Regulation by the controlled TF is measured using a synthetic target promoter driving YFP expression integrated to the *galK* locus. The target promoter is designed to be unregulated except by a single binding site for the controlled TF. The sequence and location of this binding site can be systematically controlled. (C) The quantitative regulation is measured as the fold-change in YFP expression as a function of mCherry signal. (D) The range of TF concentrations explored for each TF is shown. Here we have converted the arbitrary mCherry fluorescence signal into number of TFs using fluctuation counting methods detailed further in the SI.

In each of these individually tunable *tf-mCherry* strains, a target promoter that drives YFP expression is integrated into the genome at the *galK* locus, see Figure 2B. The basic promoter incorporates a modified *lac* RNAP binding sequence where the RNAP occupancy term, *P*, was measured previously [54]. Otherwise, the promoter is designed to be free of specific known TF binding sequences. To study the regulatory role of a specific TF, we introduce a TF-specific binding site (chosen from an array of known binding sites with strong evidence of that particular TF binding [57, 58]) cloned either directly downstream of the transcription start site (TSS), annotated as +1 for simplification or centered at 61 bases upstream of the TSS, annotated as -61. The effect of the TF binding to the promoter is then measured in terms of YFP fluorescence protein expression as a function of average number of TFs per cell for a given induction condition (Figure 2C). For this work, our focus is to select a binding site that will bind the TF. The affinity of the site or how the particular choice may influence the regulatory parameters is not something we explore exhaustively here. The specific binding sites chosen for each TF can be found in the supplemental material.

### Regulatory response of six different TFs at +1 and -61

Figure 3A-F shows the fold-change in YFP expression (promoter activity) as a function of TF copy number for the TFs examined in this study measured using single-cell fluorescent microscopy. In these plots regulation at +1 is shown as red points, regulation at -61 is shown as green points and a control promoter with no TF binding site is shown in blue. The six TFs display diverse regulatory behavior which depends on both the TF identity and TF binding location on the gene. For example, CpxR (figure 3F) is a repressor when bound at +1 but activates at −61, AscG (figure 3D) is a strong repressor at +1, however has almost no regulatory role at −61, and BetI (figure 3E) represses at both +1 and −61 but much more weakly at −61 despite binding to the same sequence at both locations. One commonality between the curves is that at +1 every TF in this study acts as a repressor. Naively the “strength” of this repression appears rather diverse; the repression from some TFs reduces YFP expression by 10^−3^ in fold-change (BetI), while others never drop below 10^−1^ (AgaR). The solid lines in this figure show a fit to the model in Figure 1A (boxed equation). In this case we are explicitly measuring the number of TFs, *N*_TF_, and we fit for the two unknown parameters *χ* and FC_max_ (see Figure 1A). In four of the six curves (Figure 3A,D-F), the +1 regulation data gives FC_max_ consistent with zero. This result is consistent with the regulatory mode of perfect repression; *i*.*e*. that the TF completely shuts the gene off when bound. As such, the difference in regulation at +1 between the TFs must be attributed entirely to either binding affinity or differing levels of stabilization (*β*) between TFs. A typical assumption is that TFs operating at +1 regulate by steric hindrance (*β* = 0), such that when the TF is bound at +1 polymerase binding is occluded. Previous studies [59, 46, 18, 60] support this assumption. The remaining curves (Figure 3B,C) show FC_max_ of order 10^−1^, however in both cases the binding is weak to the point we do not see the expected saturation of fold-change with TF copy number and, as such, the data from these curves is also consistent with FC_max_ of zero with a slightly increased value of *χ* to compensate. The collapse of all +1 regulation data to the perfect repression contour (FC_max_ = 0) is demonstrated in Figure 4A where we plot the fold-change against the effective TF concentration, *χN*_TF_ for these six TFs; the +1 data for all six TFs largely fits to a single regulatory contour associated with perfect repression once the extrinsic features such as TF copy number and binding affinity are “normalized away”.

**Figure 3:**
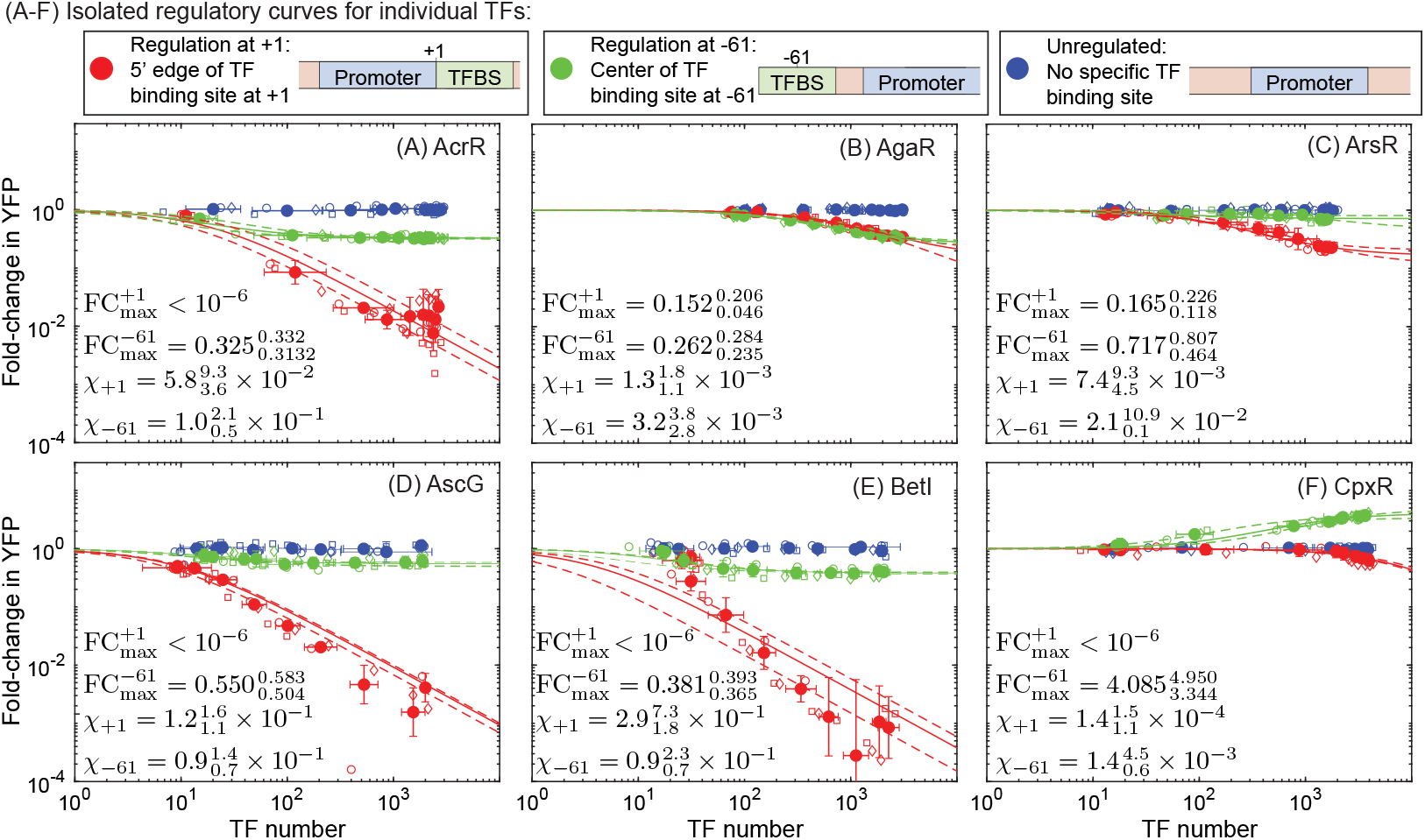
Regulatory curves for individual TFs. Each curve shows the response of a gene regulated only by the controlled TF to a measured level of TF. In all cases, the average number of TFs from each induction condition is found by converting the arbitrary fluorescence signal of each TF-FP fusion to TF number through a fluctuation counting method. For the no binding site control data (blue points), the fold-change is typically 1 for all TF concentrations; in other words, their is no regulatory response to the TF in the absence of a binding site. When the binding site is inserted just downstream at +1 (red points), the observed regulatory function is always repression. However, at -61 (green points) the response can vary between repression that is as strong as +1 (*i*.*e*. AgaR in B), repressive but weaker than at +1 (*i*.*e*. AcrR or BetI in A and E), or it can have the opposite role and activate (*i*.*e*. CpxR in F).

**Figure 4:**
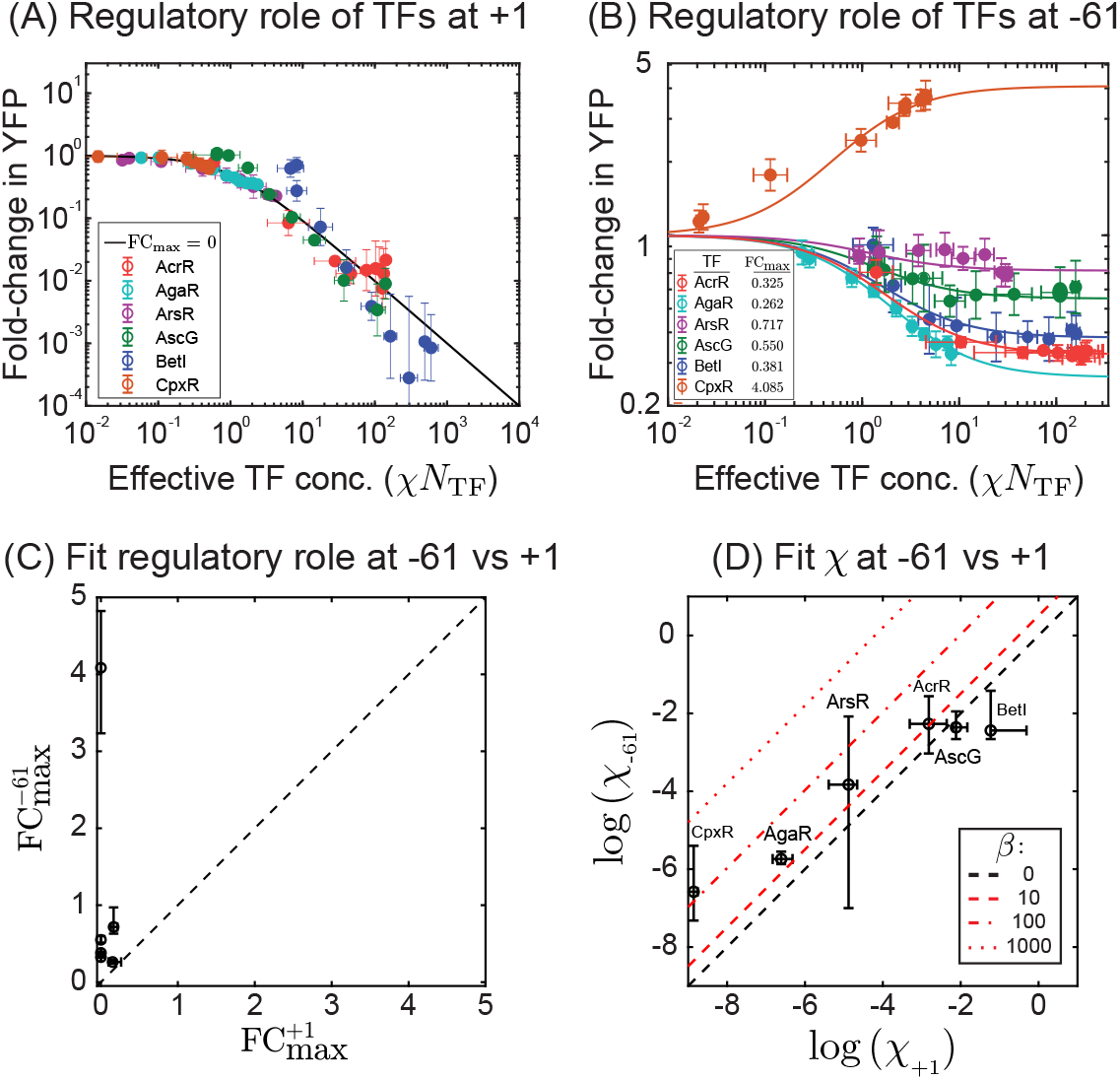
Fold-change for a given regulatory location vs *N*_eff_. The regulatory curve for all TFs when acting at (A) +1 and (B) −61 plotted against *N*_eff_ = *N*_TF_exp(−Δ*ϵ*_TF_). In all cases, the binding energy and FC_max_ are determined from fitting the equation in Figure 1A to the +1 and −61 data independently. Although the data for +1 is well described by a single regulatory behavior for every TF (pure repression, *i*.*e*. FC_max_ ≈ 0), the same TFs at -61 have a spectrum of quantitatively distinct regulatory behaviors. (C) There is no correlation for the overall regulatory role of the TF between +1 and −61, indicating position dependence for the regulatory role of these TFs.(D) The inferred TF binding affinity is consistent between +1 and −61 for all but two TFs corresponding to AgaR and CpxR, possibly indicating a contribution from TF stabilization (*Pβ >* 1).

Although at +1 the fold-change curve for each TF collapsed on a unifying regulatory profile (with FC_max_ = 0), at -61 these TFs operate with a diverse range of regulatory effects; some TFs mirror the function at +1, showing a similar profile to the response function at +1, while others show limited repressive capabilities that saturate at specific fold-change values (FC_max_), while other TFs activate expression at -61. In order to quantify FC_max_ and *χ* for each TF acting at −61, we fit the data in Figure 3 with the theory in the fold-change equation above. We find that the repressive TFs have FC_max_ values ranging between 0.2 and 0.7 while the lone activating TF is around 4. In Figure 4B, we plot the fold-change data against the effective TF concentration, *χN*_TF_. Now, rather than each TF following the same trajectory, the regulation data for these six TFs follow unique trajectories corresponding to specific values of FC_max_. Figure 4C, demonstrates that FC_max_ at +1 and −61 for these six TFs is not correlated between the two locations.

In Figure 4D we show the fit value of *χ* for each TF at −61 against the fit value of *χ* at +1. Recall that *χ* is composed of the product of two effects: the binding affinity and the stabilization. We do not expect the TF binding energy, Δ*ϵ*_TF_, to depend on where the binding site is located since the binding sequence is not changed [57]. The other possible contribution to *χ* comes from stabilization and takes the form (1 + *βP*) which means that *β* needs to be much larger than *P* for stabilization to impact *χ*. We expect the data points will fall into two potential outcomes. Points that lay on the black dashed line indicate that stabilization is not playing a significant role in the regulation at -61. However, points that are above the black dashed lines imply that the TF utilizes stabilization of RNAP (shown as red lines in Figure 4D). As can be seen, many of the data points are consistent with small or zero *β* but two TFs, corresponding to AgaR and CpxR, are significantly above this line implying that stabilization may be playing a role in their respective regulation. The implied magnitude of stabilization is shown by the red lines in the figure. Interestingly, AgaR in this case is a repressor but appears to impart strong stabilization, suggesting that even though this TF stabilizes (*β >* 1) the polymerase at the promoter it more strongly decreases the rate of transcription from polymerase bound at the promoter (*α <* 1) resulting in a net repression of gene expression. This highlights a mechanism of repression that is fundamentally distinct from the downstream (+1) position regulation for the AgaR, and demonstrates an incoherent regulatory strategy where the TF engages RNAP at two distinct steps with opposing effect.

### Profiling the spatial regulatory landscape of the CpxR TF

We now examine how the regulatory parameters that quantify stabilization (*β*) and acceleration (*α*) vary with binding location for one TF, CpxR. As demonstrated in Figure 5A, CpxR naturally binds to a wide range of promoter locations in order to regulate dozens of different genes in *E. coli*. However, the regulatory role of CpxR as a function of binding location is unclear from this data; both repressive and activating interactions are attributed to many of the locations upstream of the promoter. Furthermore, we have evidence that CpxR is capable of regulating through stabilization at −61 implying that we may be able to separate regulation through stabilization and regulation through acceleration in a more thorough examination. To measure the isolated regulatory behavior of CpxR as a function of binding location, we take the same synthetic target gene (Figure 2B) and move the TF binding site between positions centered at -48 base pairs from the TSS to -112 base pairs from the TSS (see Figure 5B). We chose this range because the vast majority of natural CpxR binding sites occur within these limits (Figure 5A). In order to enable rapid cloning and measurement, the target gene is cloned into a low-copy plasmid (rather than integrated into the genome) and fold-change in target expression and TF abundance is measured using flow cytometry (rather than single-cell microscopy). We find consistent results with the target gene on plasmid or integrated into the genome (see supplemental material).

**Figure 5:**
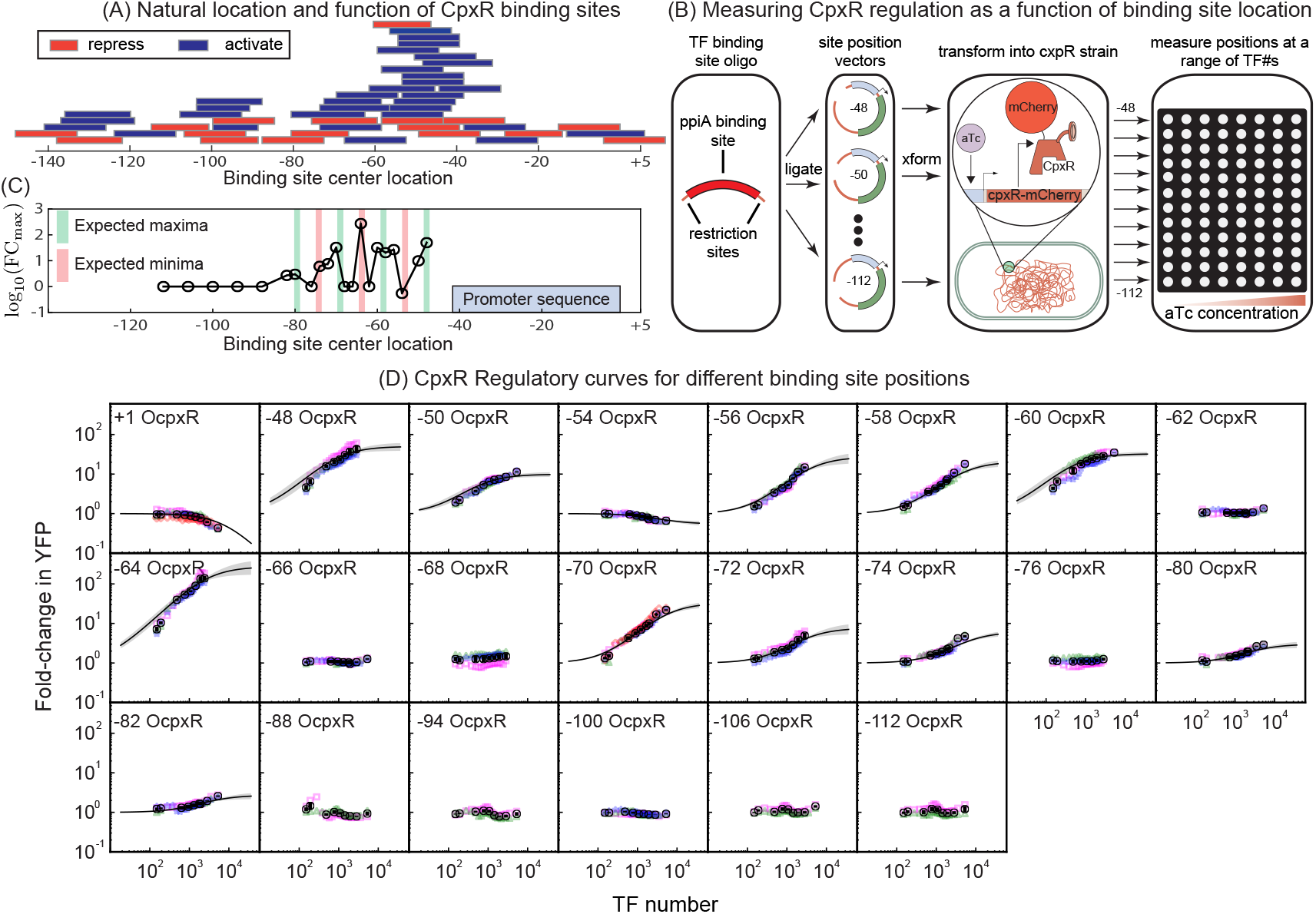
Position specific regulatory profiles of the CpxR TF. (A) The distribution of CpxR binding sites across all naturally occurring genes in *E. coli*. Note that the length of the rectangles represents the span of the binding sequence and the border color represents activation (blue) or repression (red). A majority of the binding sites are centered between the -40 to -80 positions. (B) Schematic strategy for constructing and measuring CpxR acting at a specified binding site(the *ppiA* binding sequence) inserted at 21 upstream positions and 1 downstream position on the promoter. (C) The inferred maximal fold-change (FC_max_) for each of the 21 upstream binding locations as a function of the binding location at the promoter. The center of the red and green shaded areas denote the presumed locations for the minima and maxima of the regulatory response (anchored on −48) based on the 10.5bp periodicity of B-Form DNA.(D) The regulatory profile of CpxR as a function of TF copy number for the 22 binding locations. Each panel shows the TF copy number on the *x*-axis and the fold-change in YFP on the *y*-axis of individual replicates (colored points). The black points represent the mean and standard error of these replicates. The dashed line running though the data points is the theory prediction based on inference from the model detailed in Figure 6. The shaded regions represent ±2 standard deviations of the inferred model parameters.

The data for fold-change as a function of CpxR copy number for the binding location sweep is shown in Figure 5D. Regulation of the positions studied here is primarily activation, with 11 positions showing increased expression ranging from 2-fold to over 100 fold and just one upstream position showing moderate repression. We find that the regulatory effect of CpxR depends strongly on binding site location; the −54 binding location shows weak repression but is flanked by activating positions only a few bases away (−50 and −56). As expected, positions that are far from the promoter (in this case beyond roughly 82 base pairs) show little-to-no regulation. In Figure 5C, we show fit values of FC_max_ as a function of binding location on the promoter for this data. Interestingly, we expected to see an 11 base periodicity in FC_max_ corresponding to the helicity of DNA [61, 52, 62]; we see this very roughly with maxima in activation around -48, -60, -70 and -80 (±1 bp), however -64 is a strong outlier which is expected to be close to a minimum but is strongly activating as measured here.

To extract the regulatory parameters *α* and *β* precisely from this data, it is helpful to replot the fold-Change data using a “position manifold”. Demonstrated schematically in Figure 6A, the position manifold plots the fold-change at one position against the fold-change of another position, each data point in this space corresponds to a measurement of both TF binding positions at the same TF concentration. In this case, we chose to plot all position data against the corresponding fold-change at +1; we chose +1 because based on Figure 4A we believe regulation is “pure steric hindrance” (FC_max_ = 0, *β* = 0) here. The advantage of this approach is that it enables the inference of *β* at other positions based on the curvature of the data without the need to simultaneously infer the TF binding affinity or TF copy number (see Methods for model assumptions). As seen in Figure 6B, data in the position manifold are expected to be lines emanating from (1, 1) which is the defined fold-change of both locations when the number of TFs is zero. As TF concentration is increased, the fold-change at +1 decreases towards 0 and the fold-change at the second position will either increase or decrease depending on the regulatory function at that position; as such activating positions have curves that rise as you move towards zero on the *x*-axis while repressive positions decrease. The role of *β* is clear in this formulation. When *β* is small (compared to 1*/P*) the profiles are straight lines. However, larger *β* will cause the curve to rise or fall more rapidly than linear reaching FC_max_ at higher values of the corresponding +1 fold-change. A related approach using regulatory manifolds has been applied previously [60].

**Figure 6:**
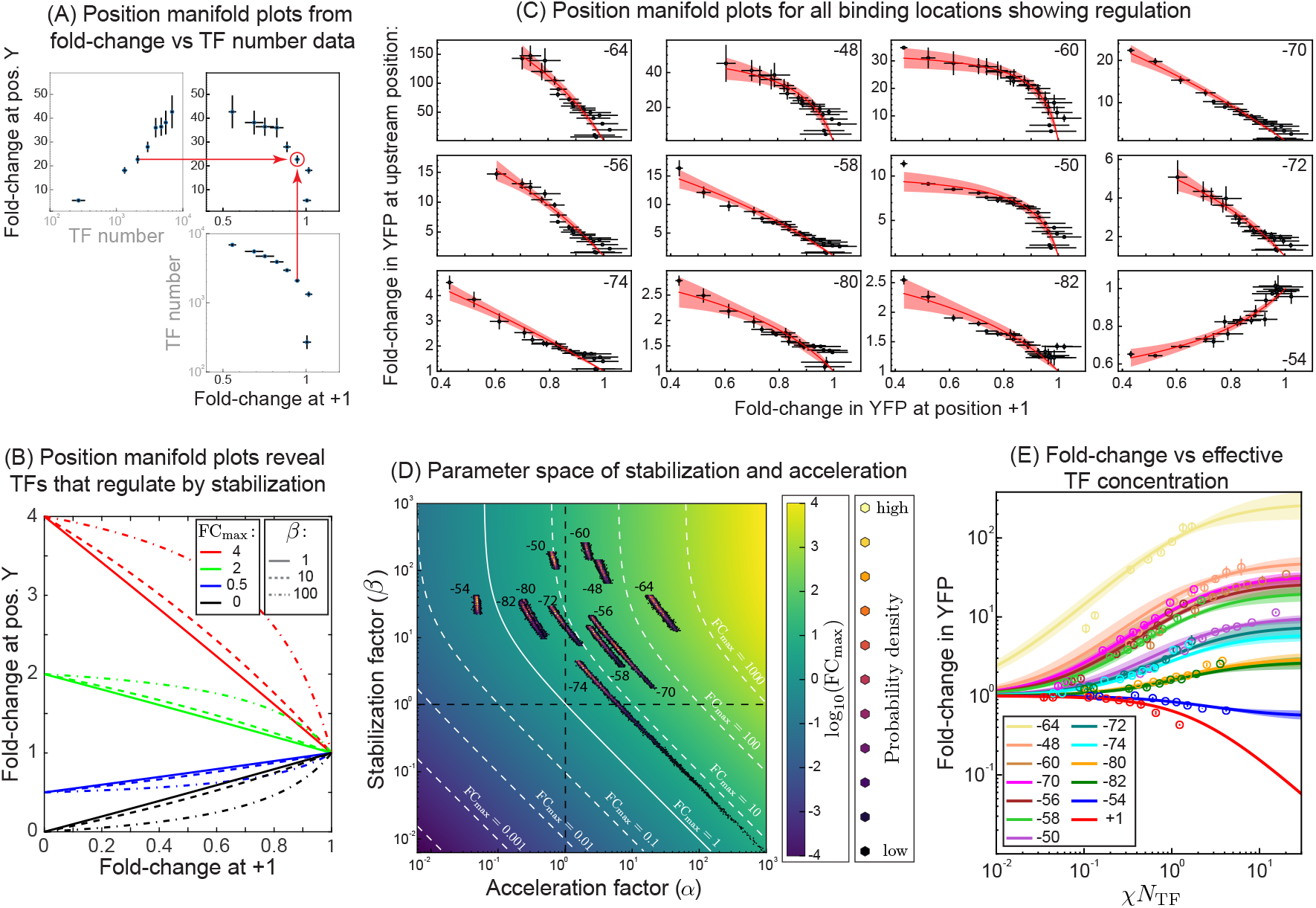
CpxR binding location determines the mode of TF-RNAP regulation. (A) Tracing out the position regulatory manifold using TF abundance from two different binding locations. (B) The position manifold curves are predicted to be straight lines when *β* is small, but curved for larger values of *β* (C) Position Manifold plots for the 12 upstream regulatory positions as a function of the fold-Change at +1. The solid red line represents the mean of the model regulatory profile generated from the inferred acceleration and stabilization parameters and the shaded regions represent ±2 standard deviations from the mean. (D) Phase plot of FC_max_ as a function of *α* and *β* parameters. The horizontal black line marks *β* = 1 (no stabilization or destabilization) and the vertical black dashed line marks *α* = 1 (no acceleration or deceleration). The white lines represent contours of constant FC_max_. The colored points represent the parameter inference of *α* and *β* for each of the regulatory positions. (E) Plot of fold-change against the effective TF concentration *χN*_TF_ for the 13 regulatory positions (12 upstream and 1 downstream) using parameters derived from the position manifold plots in (B).

In Figure 6C we replot the data from Figure 5D as fold-change at a certain position against fold-change at +1; each data point represents a measurement of fold-change at these two different binding locations for a given TF copy number within the cell. The plots are arranged from the highest FC_max_ (strong activation at −64) to the lowest FC_max_ (weak repression at −54). The solid line represents our model curve using the inferred values of *α* and *β*. Based on the inferred stabilization values, we find that the activation profiles across the position sweep is driven by varying degrees of stabilization and acceleration. The inferred values for *α* and *β* of all measured positions are shown in Table 1. We find strong stabilization in regulation at positions -50,-60, -54, -48, and -64; visual inspection of Figure 6C shows the curvature we expected to see from strong stabilization. Several positions -58, -56, -70, and -74 have regulation profiles that are approximately straight lines implying that CpxR either destabilizes or does not regulate through stabilization (*i*.*e*. (1 + *βP*) ≈ 1).

**Table 1:** Inferred acceleration and stabilization parameters. Median values of the inference chain along with the bounds that encompass the 68%th percent Bayesian credible interval of the parameters *α* and *β*. The “weak stabilization limit” limits the precision of inference estimates for positions −58, −70, and −74.

| Binding Position | $\alpha$ | $\beta$ |
| --- | --- | --- |
| −64 | 24.911<sup>31.849</sup><sub>14.911</sub> | 29.545<sup>44.169</sup><sub>12.984</sub> |
| −48 | 3.506<sup>4.008</sup><sub>2.898</sub> | 116.257<sup>150.131</sup><sub>66.453</sub> |
| −60 | 2.161<sup>2.33</sup><sub>1.974</sub> | 219.757<sup>270.481</sup><sub>151.499</sub> |
| −70 | 6.995<sup>9.748</sup><sub>3.101</sub> | 6.224<sup>9.316</sup><sub>1.776</sub> |
| −56 | 3.38<sup>4.496</sup><sub>1.745</sub> | 14.723<sup>21.647</sup><sub>5.304</sub> |
| −58 | 3.107<sup>4.127</sup><sub>1.635</sub> | 10.519<sup>15.386</sup><sub>3.796</sub> |
| −50 | 0.675<sup>0.736</sup><sub>0.612</sub> | 161.766<sup>199.140</sup><sub>108.281</sub> |
| −72 | 0.813<sup>1.053</sup><sub>0.451</sub> | 20.905<sup>30.744</sup><sub>7.678</sub> |
| −74 | 2.329<sup>3.944</sup><sub>0.467</sub> | 2.982<sup>4.685</sup><sub>4.789×10<sup>−6</sup></sub> |
| −80 | 0.289<sup>0.355</sup><sub>0.197</sub> | 27.025<sup>37.948</sup><sub>10.944</sub> |
| −82 | 0.271<sup>0.334</sup><sub>0.182</sub> | 24.811<sup>35.501</sup><sub>9.835</sub> |
| −54 | 0.049<sup>0.051</sup><sub>0.047</sub> | 34.496<sup>43.447</sup><sub>23.154</sub> |

Figure 6D shows a heat map for log (FC_max_) as a function of the regulation parameters *α* and *β*. The dashed black lines, which denote *α* and *β* equal to 1, divide the map into four quadrants, each quadrant with a specific qualitative regulatory scheme. The top-right and bottom-left quadrants represent coherent regulation strategies where *α* and *β* both contribute to activate (top-right) or repress (bottom-left) gene expression. On the other hand the top-left anad bottom-right quadrant are incoherent in the sense that *α* and *β* have opposing regulatory effects: TFs in these quadrant either slow the initiation rate of transcription while also stabilizing polymerase at the promoter (top-left) or increase the initiation rate of transcription while destabilizing the polymerases presence at the promoter (bottom-right). The solid, white contour in this plot shows where these two effects balance and the net fold-change is 1, left of this line represents TFs that repress, right of this line represents TFs that activate. Contours of constant FC_max_ are drawn as white dashed lines marking 10 fold increases/decreases in FC_max_. On this same plot we also show the inferred probability distribution of the parameters *α* and *β* for each position in our data. The black points are lower probability with lighter points representing higher probability values for the *α* and *β* parameters. One notable phenomenon is for positions with *β* less than roughly 10, the inference begins to fail for *α* and *β*. This results in inference clouds with “tails” that stretch across quadrants and precludes an assessment into the mode of regulation (see position -74). The alignment between the inference clouds of these positions and the constant FC_max_ contours, however, shows that while we make very precise estimations of FC_max_, the values of *α* and *β* are less certain and correlated. This is an unfortunate consequence of the weak stabilization limit in our model.

Interestingly, at some positions (−54, −50, −80 and −82), we see the incoherent behavior discussed above where the TF both stabilizes polymerase at the promoter and also slows the rate of initiation, essentially serving opposing functions in influencing gene expression. For −54 the net effect of these opposing mechanisms is repression, while at −80, −82 and −50 the result is activation. However, the positions with strong activation signatures (−48, −60, −64) as well as some intermediate ones (−56, −58, −70) have stabilization and acceleration values that impart a coherent strategy of regulation where RNAP recruitment and acceleration of transcription work together. Interestingly, all but one of the regulatory positions studied here show clear positive stabilization (*β >* 1), even the lone repressive position (−54). In our data, upstream regulation by CpxR typically involves stabilizing RNAP, regardless of the net regulatory function (repression or activation). However, the level of acceleration/deceleration varies more significantly between positions from a roughly 20 fold deceleration, which results in overall repression of expression, up to a 25-fold acceleration, which results in strong activation.

Finally, combining the inferred regulatory parameters, *α* and *β* determined through inference in the position manifold space, with the measured extrinsic features (TF copy number and binding affinity) of gene regulation produces model curves using the effective parameterization that fits our data well. The effective parameters FC_max_ and *χ* for the CpxR TF data (See Methods section for details) are used to plot the fold-Change as a function of *χN*_TF_ for the 13 regulatory positions in Figure 6E. Crucially, we demonstrate that the two key variables *χ* and FC_max_ are effective in capturing the fold-change position sweep profiles similar to the plot in Figure 4A and B.

## Discussion

To build a predictive understanding of gene regulation, we need to understand not just where and when TFs bind but also learn the function and magnitude of the mechanisms of regulation at work by each TF [63, 64, 65]. Often, it is difficult to separate the contributing factors of regulation, such as the TF binding affinity, copy number and interactions with other TFs, from the regulatory role of the TF that is characterized by its interactions with RNA polymerase at the promoter [66]. Here, we use a synthetic biology approach to measure the isolated regulatory effect of a TF on an otherwise constitutive promoter. This data is interpreted through a thermodynamic model of gene expression that treats the regulatory role of TFs as a combination of interactions that stabilize (or destabilize) the polymerase at the promoter and interactions that accelerate (or decelerate) the rate of transcription when the TF is cobound with polymerase. The model used here allows for both modes simultaneously and, importantly, enables us to quantify TF regulatory function continuously rather than categorically as “activators” or “repressors”. Using this model, we are able to characterize the huge range of regulation we see from the TFs in this study, which ranges from 10,000 fold repression up to 100 fold activation and everything in between with the same model. We believe this fluid classification of TF function can be a useful tool for characterizing TFs for the purpose of model building and predictive design of gene regulation.

We found that for TFs operating immediately downstream of the promoter, the regulation of each TF was consistent with strong repression (*i*.*e*. with FC_max_≈ 0). Despite the large range in magnitude of regulation at this location, the same intrinsic regulatory mechanism seems to be conserved; differences in the magnitude of regulation was primarily due to difference in TF binding affinities rather than in the fundamental regulatory mechanisms of the TFs. In contrast, when these same TFs bind 61 base pairs upstream of the promoter the regulatory function of the TFs varied more substantially. We find some TFs remained as strong repressors (similar to their function at +1) however other TFs only weakly repress expression regardless of TF copy number. We attribute this to intrinsic properties of the TF, polymerase (de)stabilization and (de)acceleration of transcription initiation by the TF, which depend on TF identity. Furthermore, profiling the regulation of CpxR at upstream binding locations, we find that this TF is capable of regulating multiple steps of the transcriptional process simultaneously and joins a growing body of evidence for TFs engaging in complex regulatory maneuvers at the promoter [67, 68, 69]. This insight into how activation is actually brought about by the independent contributions of acceleration and stabilization demonstrates the applicability of the model to *in vivo* data, where previous work analyzing the kinetics of activator regulation have relied on *in vitro* biochemical assays [70]. We find that the contributions of these two mechanisms do not correlate with position; in some locations we found stabilization and acceleration acting together to produce strong activation, in other locations deceleration and stabilization worked incoherently resulting in weaker activation and repression. A startling feature of this incoherent regulation was its presence in the regulatory response of both an activator (CpxR) and a repressor (AgaR) and demonstrates that this type of regulation is not just accessible to TFs, but may be a pervasive aspect of TF-promoter regulation.

The concept of stablilization and acceleration working antagonistically, with the step carrying the larger effect size ultimately determining the status of expression (activation or repression), has implications for the current paradigm of viewing TF regulation, particularly activation in the context of Class I and Class II promoters. This delineation of promoter class is based on the type of molecular contacts the activator makes with RNAP [71, 19, 72, 73], and addressing how these contacts shapes the relative effects of *α* and *β* would fill a vital gap in elucidating the molecular determinants that give rise to either coherent and incoherent regulatory modes. Such information would allow for a more complete characterization of TFs, and in conjunction with methods of profiling TFs through genome wide occupancy techniques provides an important edge in the challenge of uncovering an “expression” code: i.e, a set of rules that govern the magnitude and duration of gene expression from natural promoters [74, 75, 76]. Realizing this goal, however, requires a systematic characterization of the TF position-regulatory axis to determine the spatial landscape of *α* and *β* and provides a compelling basis for characterizing a broader set of TFs to generate a “regulatory compendium” where TFs are classified according to their regulatory mode. This argument, however, is predicated on our ability to infer regulation driven by stabilization and acceleration with sufficient precision. One unfortunate feature of our experiments, as designed, is our inability to measure destabilization with good precision (*β* ≤ 1) and limits the characterization of TFs that engage through this mode. One potential way to overcome this is to select a stronger promoter sequence for the target gene. The selected sequence we used in our synthetic circuit was an attenuated form of the *lacUV5* promoter, which was selected to increase the dynamic range of the promoter. As such, measuring the isolated regulatory function of TFs may require a range of promoters to fully characterize the wide array of regulatory behaviors possible.

Here we focus entirely on the isolated regulatory role of each TF, however it is clear that one of the next steps is to probe how quantified TFs regulate together. the ability to distinguish and quantify the stabilization and acceleration mode of regulation and characterize TFs based on them is important for developing general theories of regulation that include TFs acting on different kinetic steps of the transcription process [77, 78, 79]; predictions for the combined regulatory effect of two stabilizing TFs should be different than predictions for a stabilizing TF acting together with an accelerating TF [77]. With each TF characterized, we can develop an empirical baseline or null hypothesis for what a TF should do on a gene: departures from this expectation, due to the emergence of complex regulatory phenomenon brought about by TF-TF interactions [63], allosteric interactions [80, 81] or other effects indicate surprises that warrant testing in these expanded models.

## Methods

### Thermodynamic model for single TF regulation

In the work here we use the standard form of the thermodynamic model as derived elsewhere [46, 48, 50]. In our framework, we include an additional consideration of the TF altering the rate of transcription as detailed in [52]. The partition function for this system is:

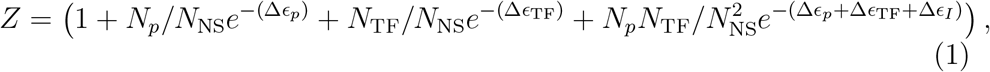

where each term (in order) represents the weight of the unbound, bound by polymerase, bound by TF and cobound state. The terms *N*_*p*_ and *N*_TF_ are the total number of polymerase or TF molecules, with the term *N*_NS_ in the denominator scaling the respective terms with the total number of potential binding sites on the genome to give an effective concentration on the chromosome. The energy terms Δ*ϵ*_*p*_ and Δ*ϵ*_TF_ are the binding affinities of polymerase or TF to their promoter or operator site. The stabilization term (*β*) as discussed in Figure 1 is represented by the exponentiation of Δ*ϵ*_*I*_. The probability to find polymerase bound as a function of TF number is then,

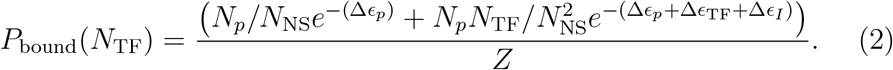

To compare with experimental measurements, we model YFP expression from our synthetic gene circuit as the convolution of the state specific transcription rates and the states enumerated in *P*_bound_(*N*_TF_). For the state in which RNAP is solely bound, we give a rate of expression as r (a course-grained parameter representing the basal rate of YFP production). For the TF-RNAP co-bound state, we assign a scaling factor *α* that represents the change in transcription rate when the TF is bound (acceleration):

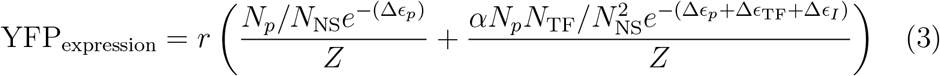

In practice, what we seek to model is the fold-change in gene expression, which is the change in expression level relative to the unregulated gene. Based on the partition function and the state specific transcription rates, the fold-change in expression then assumes the following form:

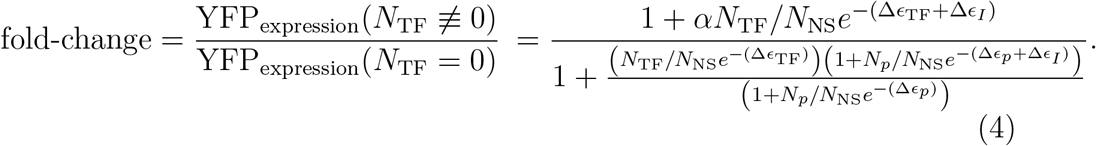

We then define the following terms 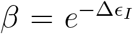 and 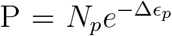. As we have measured the value of P in our synthetic circuit to to be 6.6 × 10^−2^ [54], we safely assume the weak promoter limit simplifies the expression 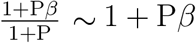. The fold-change is then written as:

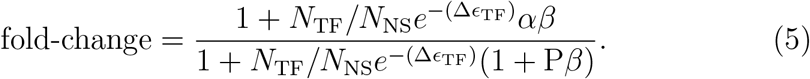

We now define the final term in our derivation: 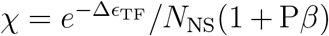 which represents the effective component that modifies the TF copy number with the product *N*_TF_*χ* acting in our model as the effective TF concentration. We now re-write the fold-change in terms of the effective TF number (*χ*N_TF_), and the maximal fold-change 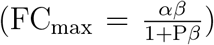 as presented in Fig1 and the main text.

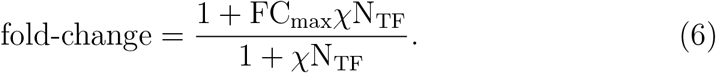

### Choice of the core-promoter sequence used in the synthetic circuit

Previously we have found that the weak promoter approximation describes *in vivo* measurements of repression of the *lac*UV5 promoter by LacI [21, 18]. For this study, where we expect to find both activation and repression, we decided to use a weaker promoter for the target gene. This promoter was designed such that 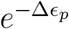 is roughly 1*k*_B_*T* lower than that of *lac*UV5 [54]. We have confirmed that the basal expression of this promoter decreases as expected and that regulation follows the same quantitative response to LacI as for *lac*UV5 [54]. Given that we have previously measured the core-promoter strength used in our synthetic circuit to be at *P* = 6.6× 10^−2^ [54], we expect that the approximation 1 + *P*≈ 1 used in deriving the thermodynamic predictions for the fold-change in gene regulation is justified in our work. The choice of the promoter sequence comes with its trade-offs: a weaker promoter sequence, while allowing for a larger window of measurement for activation, potentially limits the ability to measure smaller *β* values for activation (the weaker the promoter, the larger the range of beta that is constrained to measurable dependence with *α* - note the weak stabilization limit discussed in the Main Text). Taking all points into consideration, we feel that our choice of the promoter sequence adequately balances competing objectives of detecting activation, and inferring the contributions of *α* and *β*, and allowing for strong enough expression to measure 100 - 1000 fold repression at +1.

### Position Manifold Derivation

As our primary motivation in this study rests on changing the binding location to explore the regulatory properties of a particular TF, we looked for a way to reformulate the thermodynamic model in such a way as to remove the binding affinity parameter from our consideration (under the assumption that the binding affinity is set primarily by the TF binding sequence which is invariant). This would allow us to infer the intrinsic regulatory features for a TF (the acceleration and stabilization parameters). To do this, consider the following reformulation of the thermodynamic model where the TF concentration is written as a function of the fold-change (abbreviated FC below) for a given binding regulatory location (designated by the superscript x or y):

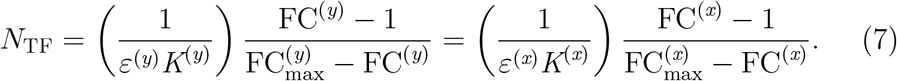

Given that we measure TF abundance, we can essentially couple the fold-change in regulation between two different positions by allowing the TF concentration to trace out a manifold that specifies the fold-change at positions y as a function of the fold-change at position x.

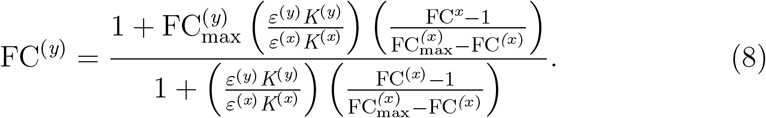

As in the first section of the Methods, we define 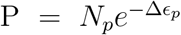 and 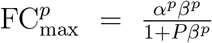 where the superscript *p* represents the regulatory position. We also introduce two new terms for compactness 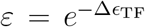 and 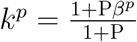. Assuming the binding affinity is constant between the two locations (*ε* = *ε*^*x*^ = *ε*^*y*^) we achieve a reduction in the manifold:

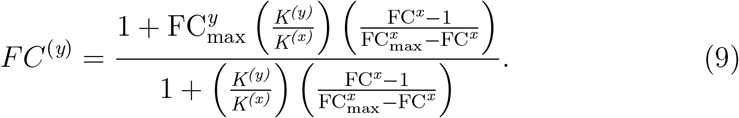

Based on this reformulation, the key parameters to consider are the acceleration parameters (couched in the FC_max_ term as described in Fig1) and the stabilization parameters at positions x and y (a total of 4 parameters). In a sense, the “position manifold” allows us to remove what we consider to be the extrinsic feature of TF regulation (the binding affinity and TF copy number) from the intrinsic features of the TF regulatory response (the regulatory activity as of the TF on RNAP through stabilization and acceleration). We can further simplify the model by taking into account a judicious binding location for the position x. Taking the +1 binding location, where the assumption of steric hindrance in our thermodynamic model for all the TFs surveyed in this work is justified, we set *β*^*x*^ ∼ 0 (which makes *k* = 1) and 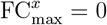. This leads us to the final form of the position manifold for a given regulatory position (as a function of the fold-change at the +1 position - FC^+1^):

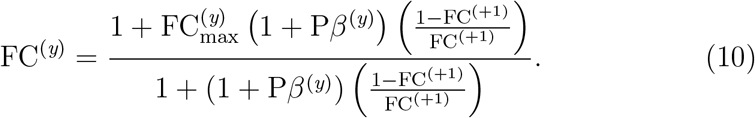

Now we see that the only parameters that remain in the model are the acceleration and stabilization at position y (the regulatory position under consideration).

### Culture conditions and Data acquisition procedures for Microscopy and Flow Cytometry Data

The position dependent regulatory profiles for the 6 TFs -AcrR, AgaR, ArsR, AscG, BetI, CpxR - evaluated at +1 and −61 positions were measured using fluorescence microscopy. At every microscopy session, the TF titration strains (harboring the integrated TF-mCherry fusions in the *ybcN* locus and the synthetic circuit in the *galK* locus) were cultured with companion strains. These include the TF-mCherry fusions lacking the *galK* synthetic circuit integration (necessary to derive the calibration factor to convert the arbitrary fluorescence signal into TF copy number) and TF-mCherry fusion strains with the TF binding site missing from the integrated synthetic circuit (necessary to account for TF titration effect on gene expression). Further-more, constitutive strains lacking the TF-mCherry fusion (integration in the *ybcN* locus) and expressing the integrated synthetic circuit were necessary to compute the fold-change in gene expression.

Single colonies of bacterial cultures from freshly streaked LB-Agar plates with appropriate antibiotics are grown overnight in 1 mL of LB in a 37° C incubator shaking at 250 rpm. Cultures are diluted 10^4^ −10^5^ fold into 1 mL of fresh M9 minimal media supplemented with 0.5 percent of glucose at different aTc concentrations (0, 0.25, 0.5, 1, 1.5, 2, 3, 4, 5, 8, and 10 ng/mL) and allowed to grow at 37° C until they reach an OD600 of 0.1 to 0.4 and harvested for microscopy. 1 *µ*L of cells is spotted on a 2 percent low melting agarose pad (Invitrogen 16520050) made with 1X PBS. An automated fluorescent microscope (Nikon TI-E) with a heating chamber set at 37°C is used to record multiple fields per sample (between 6-12 unique fields of view) resulting in roughly 100 to 500 individual cells per sample.

The calibration factor for the conversion of mCherry fluorescence to TF copy number is quantified by measuring the fluctuations in fluorescence partitioning during cell division [18]. Briefly, cells expressing the TF-mCherry fusion protein are grown as described above, and just before imaging 100 *µ*L of cells from different aTc concentrations are pooled together and washed twice with M9-glucose minimal media containing no aTc. Cells are then spotted on 2% low melting agarose pad made with M9-glucose minimal media. Phase images are captured for roughly 150 to 200 fields and their positions are saved for later. These phase images (named as Lineage tracker) will serve as a source file for lineage tracking of the mother-daughter pair. After one doubling time (roughly 1 hour or depending on the doubling time for different TF strains), the microscope stage was returned to the same field of view using the saved position matrix and are imaged again (and named as daughter finder) using both phase and mCherry channels

To measure the regulatory profiles for the CpxR TF position sweep constructs, we used flow cytometry for rapid and reproducible data acquisition. The CpxR-titration strains harboring the position regulation plasmids (See Strains) were cultured in LB + Kanamycin media from single colony inoculates until saturation. A 1:10000 dilution for each strain was then made in M9 minimal media supplemented with glucose along with the appropriate amount of aTc to titrate CpxR-mCherry levels. We found that the following aTc concentrations 0, 0.5, 1, 1.5, 3, 4, 6 and 8 ng/ml provided a good dynamic range of TF expression while maintaining viability of the CpxR strains.

The aTc dilution solutions were made from a stock solution of 1 mg/mL suspended in ethanol and were made fresh prior to the application of the aTc for the M9 culture. After M9 dilution, the strains were grown in 96 well plates to steady state (OD600 of 0.1 – 0.2). Similar to microscopy acquisition procedure, we had constitutive (CpxR-KO) strains transformed with the binding position plasmids along with the CpxR-titration strain transformed with the plasmid having the TF binding sequence removed from the promoter to account for physiological effects of CpxR-mCherry titration to calculate the fold-Change. Cells were diluted between 1:16 to 1:32 fold in PBS media in a 96 well cytometer plate prior to data acquisition and cytometry was performed on a MacsQuant VYB. At the beginning of each run, an initial gating strategy involving the FSC and SSC height information was used to eliminate background events and samples were run to achieve ∼60000 gated events for each position strain at a given aTc concentration.

### Data processing steps for Microscopy and Cytomtery Data

For the +1 and −61 position sweep regulatory curves of the 6 TFs, we took the microscopy images and segmented individual cells using a modified version of the Matlab code Schnitzcells [82].We use this code to segment the phase images of each sample to identify single cells. Mean pixel intensities of YFP and mCherry signals are extracted from the segmented phase mask for each cell. The autofluorescence is calculated by averaging the mean intensity of the autofluorescence strain in both mCherry and yfp channels and is subtracted from each measured YFP or mCherry value. Total fluorescence for each channel is obtained by multiplying the mean pixel-intensity with the area of the cell. Fold-change in expression for a given binding site is calculated by the ratio of total fluorescence of strains expressing the TF to the strains with no TF. For partitioning statistics to estimate the calibration factor, mother-daughter pairs are first automatically identified and verified manually to ensure cells made exactly one division. The mean pixel intensity and area of the mother-daughter pairs are obtained. The background fluorescence is estimated as described in Ali *et al* [7] using the inverse mask of individual frames. The sum and difference in fluorescence of the two daughters were then used to find the conversion factor, *v*, between fluorescence and number of TFs using the equation (*I*_1_−*I*_2_)^2^ = *v*(*I*_1_ + *I*_2_), which stems from the assumption of binomial partitioning of TFs at cell division [82].

For the CpxR position cytometry data, we adapted a robust data analysis procedure [83] to computationally gate events to ensure reproducible fold-change measurements for a given position across replicates. For a given position strain replicate measurement, we collected the data across all the aTC concentrations and proportionally binned the single flow cytometry events into 16 RFP intervals (intervals were off unequal size in RFP space with the number of cells in each bin more or less constant). We then took these binned evensts and gated them using the log_10_ values of the Forward Scatter and Side Scatter height profiles for each event (referred to as FSC and SSC respectively) to improve the likelihood that the final retained events were single cell measurements. To construct this gate, we computed the mean and covariance matrix for eact data set for every RFP bin and used these statistics to fit an ellipsoid to the full data set according to the following formulae:

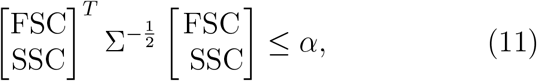

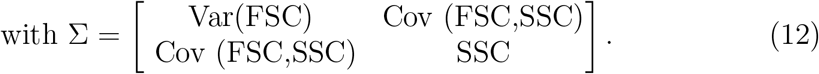

This step retains events that are within a particular distance from the center of the ellipsoid using an appropriate value for the cut-off (alpha). We based the value of the cut-off on the following rationale: As each event is essentially a vector of FSC and SSC values (the log_10_ values of the Forward Scatter and Side Scatter), we assume the joint values are normally distributed, which translates to a distance metric that is a chi-squared random variable (the sum of two normally distributed entities is chi-squared with df 2). We selected *α* as the 5^th^ percentile of values from the cumulative distribution, and events within the cutoff were taken to be single cell measurements used to compute the fold-change values presented in the results section. The resulting gated events for each of the 16 RFP intervals were pooled from all position strains and replicates, and we excluded events with RFP measurements below a certain threshold determined by visually assessing the the Fold-Change profile of the “control” (*DelBS*) circuit. The YFP signal for these events had large fluctuations and we reasoned to the flow-cytometry approach probably fails in measuring cells with low TF copy number. The retained events were then binned proportionally into 22 intervals, and the median RFP and Fold-Change values for each interval was reported as the representative measurements for that bin. The choice of intervals at this step (with the exception for very large bin sizes) does not seem to appreciably alter the main findings from our inference into the acceleration and stabilization for the CpxR binding locations (see SI Figures 5 and 6 for details).

### Parameter fitting and Inference for position dependent fold-change regulation data

We interpret the promoter regulatory data from the 6 TFs surveyed at the +1 and − 61 binding locations through the thermodynamic model as specified in the results section. The fold-Change data for the TF-position strain (See Data processing steps for Microscopy and Cytomtery Data section for details) as a function of N_TF_ was fit to equation 6 with the aim of extracting the best-fit value of the FC_max_ and *χ* (the product of the stabilization effect and the TF binding affinity). We used a bootstrapping procedure to generate confidence intervals for the both the FC_max_ and binding affinity parameters, and for each of the TFs surveyed we fit the +1 and −61 parameter sets independently. The bootstrapping procedure resampled the data points from the fold-change vs RFP curve for a given TF-binding location across all replicate datasets 1000000 times. For each iteration, fold-change replicate data points from a given induction conduction were sampled to generate a possible regulatory response as a function of TF-copy number. Each of these resampled curves were then fit to the thermodynamic model outlined in Figure 1A using a non-linear least squares fitting procedure to determine the optimal fit for the values of FC_max_ and *χ* parameters. As seen in Figures 4C and Figures 4D, we report the means and confidence intervals for these two parameters and plot the curve generated by taking the expected value of the thermodynamic model conditioned on the model parameters along with the 94th percent confidence interval.

For the CpxR position sweep data, we used the position manifold formalism to delineate the values of the acceleration and stabilization parameters. For positions that showed discernible regulation (12 out of the 22 upstream positions profiled), we assume that the binding affinity is constant between the regulatory positions as the only changing variable is the binding location (the binding sequence is the same) and recast the binned data from the FC vs RFP replicates using the position manifold formalism as detailed in the Methods (see Position Manifold Derivation). To sample the probability space of the acceleration and stabilization parameters for a given binding location, we started by inferring the joint posterior distribution FC_max_ and *K* = 1 + *Pβ* using a Bayesian approach that relates the parameters underlying our thermodynamic model to the data according to the following relation:

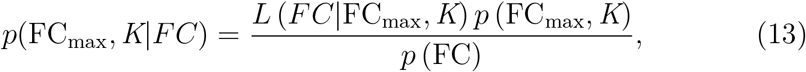

where the term on the left hand side is the posterior distribution of the parameters (FC_max_ and *K*). The terms on the right hand side represent the likelihood of the data given the parameters(*L* (*FC*|FC_max_, *K*), the prior distribution of the parameters (*p* (FC_max_, *K*)), and lastly the distribution of the data itself *p* (FC). Each of the 12 regulatory positions were fit separately using the Bayesian inference procedure, we specified our likelihood function as a normal distribution of the form:

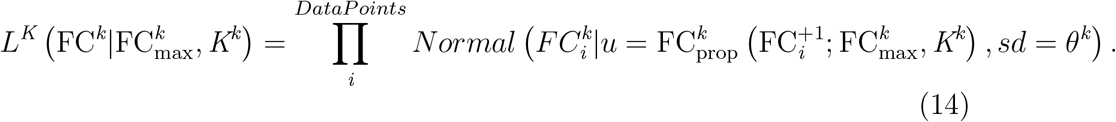

The superscript k represents an upstream binding location for the CpxR TF and the subscript i represents the data points for a given position regulatory dataset. The proposed fold-change value FC_prop_ from the model takes the form:

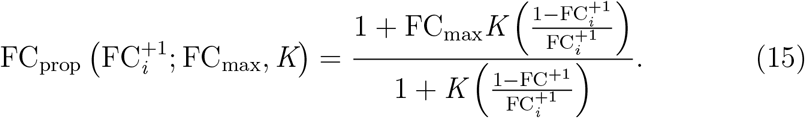

The crux of the Bayesian approach to model inference is to simulate candidate draws of the joint posterior distribution of the FC_max_ and *K* parameters by proposing candidate values from the prior distribution, generating the thermodynamic curve, and evaluating the likelihood function. A transition in the jointly sampled parameter space from one set of parameter values to another is based on the premise that parameter sets will be sampled in proportion to the probability of the posterior distribution (as long as the sampling chain is drawing from the stationary distribution). This process is repeated until a given number of draws have been made from the joint posterior distribution. The results of this inference procedure are used to draw the model curves in Figure 5C and we use the relation between the sampled parameters (FC_max_ and *K*) and the acceleration and stabilization parameters as detailed previously in the methods section. The sampling procedure was implemented with the PyMC3 package that utilizes the NUTS sampler, a particular implementation of the Hamiltonian Monte-Carlo algorithm, to sample the joint posterior distribution [84].

We used a uniform distribution as the priors for both the FC_max_ and *K* model parameters with appropriate bounds for each parameter. For *K*, we ensured that the lowest potential value is 1, in line with the assumptions from the derivation of the position manifold formalism. We checked the inferences from each position to ensure that the bounds we enforced on both parameters were appropriate and that the sampled values were not tending towards the edge of the sample space. Furthermore, we cast the *σ* parameter (the standard deviation) of the position specific likelihood function as a hyper-parameter in our sampling procedure and set the prior distribution as uniform over a defined interval with the lower bound = 0. Overall, our inference approach allowed us to ensure precise inference of *α* and *β* and for 8 out of the 12 regulatory positions (See Table 1).

To get the stabilization values from this inference procedure, we simply used the following relation between the *K* and *β* and the fact that the polymerase occupancy has a measured value of 6.65 × 10<^10−2^ in our synthetic promoter,

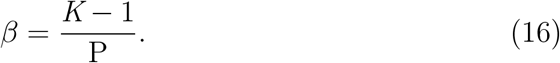

Given the stabilization value for a draw in the chain, we then find the corresponding acceleration value using the jointly sampled FC_max_ value and the relation:

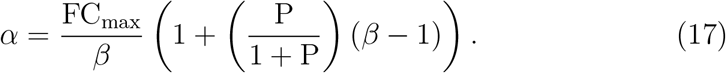

Table 1 lists the inferred acceleration and stabilization parameters. We report the median values of the inference chain along with the bounds that encompass the 68%th percent Bayesian credible interval of the parameters. For positions −58, −70, and −74, the inference estimates are not precise due to the phenomenon of the “weak stabilization” limit as discussed in the Main text. Examining the posterior distribution of the FC_max_ and *K* parameters, we find the values of the 68% credible interval for these positions (both *α* and *β*) to encompass both coherent and incoherent regimes.

### Using the position manifold parameters to generate the thermodynamic model in FC vs *N*_*TF*_ space

The parameters inferred using the position manifold approach were used to construct the thermodynamic model curves in Figure 5C and Figure 6E. To accomplish this, we used the *K* = 1 + *Pβ* and FC_max_ values inferred for each of the 12 regulatory positions (see Methods Section: Parameter fitting and Inference for position dependent fold-change regulation data). To transform the Markov chain of *K* values to the *χ* for the main thermodynamic model (Figure 1, and Methods) requires the binding affinity of the TF (the final remaining term in *χ*), and we inferred the binding affinity from the datasets encompassing all 13 regulatory positions for the CpxR TF (the 12 upstream regulatory positions and the one immediate downstream positions - the +1 position). The binding affinity was treated as a global parameter in our Bayesian inference scheme and was inferred according to the following model:

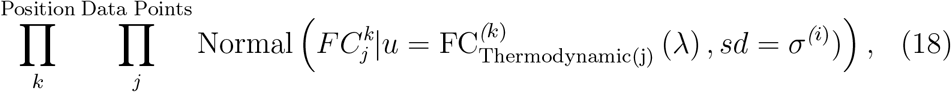

with the mean of the likelihood function specified by the thermodynamic model outlined in section: “Thermodynamic model for single TF regulation” and takes the following form:

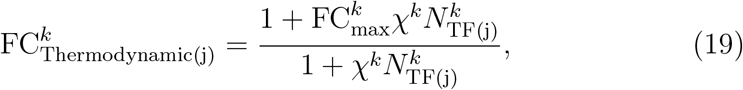

Where *χ*^*k*^ = *λK*^*k*^.The parameter *λ*, is the global parameter representing the binding affinity of CpxR TF (assumed to be constant across the regulatory positions assessed in this work). FC_*k*_ is the general thermodyanmi model specified in equation 6 with the object of our inference to infer 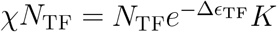. For the 12 upstream regulatory positions, as the chains of FC_max_ and *K* were available from the inference of the position manifold dataset, we inferred the global *λ* parameter to the datasets for each of those positions with the values of these two parameters determined from the mean of their respective Markov chains. For the +1 dataset, the parameter FC_max_ was set to 0 as determined in 4A, and the *K* was set to 1 in keeping with the assumption of steric hindrance. The mean value of the Markov chain *λ* inferred from this model, along with the scaling factor (see SI section - **Converting the mCherry signal to TF copy number**) was used to generate the thermodynamic model curves (mean ±2*σ*) as seen in Figure 5C and Figure 6E.

### Cloning the TF specific binding location gene circuits

The upstream promoter sequence in our synthetic gene circuit was derived from the *P*_*DL*5_ plasmid that contains a modified version of the *lacUV5* promoter sequence as used previously [18]. Binding sequences for AcrR, AgaR, ArsR, AscG, BetI, CpxR TFs were cloned at the +1 and the −61 locations of that plasmid by ordering the binding sites as a primer dimer with overhangs that amplified the surrounding flanking regions of the *P*_*DL*5_ vector where we wished to insert the sequences and PCR amplified the desired regions. This amplified sequence contained the YFP coding sequence, transcriptional termination sequence, and the 250 bp upstream promoter sequence containing the TF-biding site either at +1 or −61 along with the Kanamycin selection cassette. The amplification produced two fragments and Gibson assembly was then used to stitch the two fragments resulting in a sequence that had the desired regulatory sequence, promoter, reporter cassette along with overhangs for the *galK* locus. The resulting reaction was purified, and transformed into the *galK* locus in using the *pKM208* recombination *E*.*coli* strains [85]. The strains were then Sanger sequenced and verified to contain the appropriate regulatory and promoter sequences and used P1 transduction to transfer the *galK* locus into the approrpiate TF titration strains (both the TF knockout strain and the TF-mCherry strains).

To clone the synthetic target promoters for the CpxR position sweep regulation experiments, we designed an approach to make fast and efficient cloning of any TF binding sequence at defined locations ranging from +1 to -112bp relative to the TSS on the unregulated DNA circuit (*P*_*DL*5_). We designed forward and reverse primers to amplify the plasmid at defined locations in the promoter sequence. These primers were used to insert a *ccdb* cassette at the precise location upstream of the gene circuit and had overhangs for the typeIIs BbsI restrction site. This allowed for excision of the *ccdB* cassette and cloning of BbsI digested TF binding sites that had complementary overhangs to excised region. The *P*_*DL*5_−*ccdB* plasmids were assembled and transformed into *Escherichia coli* DB3.1 strain that harbors key mutations in DNA gyrase that tolerates the *ccdB* toxin [86] for stock cataloguing and sequencing. The plasmids were then incubated with double stranded oligos that had the TF binding sites flanked with the Bbs1 restriction sites and the 4 bp complementary sequence to the digested plasmid. This approach allowed the incubation to be a one-step Digestion and Ligation reaction to excise the *ccdB* cassette and insert the TF binding sequence at the defined locus. Stock curation, sequencing, and final transformation into the relevant stains were done as for the other TF-position constructs. All primers and TF binding sequences used for the cloning of the synthetic target genes are listed in the Supplement Section.

### Engineering the titratable TF-mCherry fusion strains

All strains used in this study are constructed from the parent strain *E. coli* MG1655. The TFs investigated in this study include AcrR, AgaR, ArsR, AscG, BetI, and CpxR. Each TF gene is deleted from its wildtype locus and expressed from the *ybcN* locus under the regulation of a Ptet promoter. The autofluorescence strain for each experiment is *E. coli* MG1655 with the corresponding TF knocked out from the wildtype locus.

We used the KEIO library [87] as the starting point for the construction of the 6 TFs with the titratable TF-mCherry fusion cassette. We selected the corresponding clone from the KEIO library with the TF-knockout (the coding and upstream regions of the TF gene are replaced by a constitutive promoter expressing Kanamycin), and deletion of TF gene from the wildtype locus was performed by P1 transduction of the corresponding knockout from the KEIO collection to the MG1655 *E. coli* strain. The kanamycin cassette was cured using the *frt* flippase expressed from *pCP20* plasmid. The TF ORF(open reading frame) was fused to mCherry with an AEK linker sequence by PCR amplifying the ORF from the MG1655 strain and cloning into the plasmid pZs3 containing the Ptet promoter with the AEK-mCherry cassette down-stream. The Ptet TF-mCherry fusion construct was then amplified by *ybcN* integration primers for chromosomal insertion. The target gene for integration is constructed by SOEing PCR using the primer pairs listed in the Supplemental Section. Chromosomal integration of the TF and target are performed by lambda red recombineering assisted by plasmid *pKM208* as described in [85]. The strain we used for the integration of the TF-mCherry fusion cassette had the inducer construct (constitutive promoter expressing TetR) already integrated at the gspI locus. After integration, we sequenced the regions surrounding the TetR promoter and the TF-Linker-mCherry cassette to confirm the regions were free of any mutations. We then catalogued and stored these strains before making the synthetic circuit measurements as presented in the Main Text.

