## Supplemental materials for "Quantifying the regulatory role of individual transcription factors in *Escherichia coli*"

### Supplementary Info

##### Measurement of TF abundance in Flow Cytometry experiments: Converting the mCherry signal to TF copy number

To measure the regulatory response of the CpxR regulated promoter at 22 binding locations, we used flow-cytometry to measure TF abundance and target gene expression (promoter activity) for cells at steady state. Our goal was to look at the regulatory level of the target gene as a function of TF copy number [1], which required TF abundance measurements to be converted from arbitrary fluorescent units to copy number of the TF-mCherry fusion molecules in the cell. To convert the arbitrary mCherry signal from the flow cytometer to TF copy number, we took the flow cytometer measurements of the Fold-Change response curve as a function of mCherry levels at the +1 position and compared it to the measurements we made using microscopy (see Figure S1). As we show in the Main Text this regulation is consistent with steric hindrance ($FC_{\max} = 0$, $\beta = 0$ [2]). We then extracted the parameter, $\chi$, from both the microscopy and cytometry curves using the thermodynamic model we present in the Main text and compared them. Details on this approach can be found in the Methods Sections **Parameter fitting and Inference for position dependent fold-change regulation data** and **Using the position manifold parameters to generate the thermodynamic model in FC vs $N_{TF}$ space**.

We reasoned that the while the value inferred for the cytometry curve ($\chi_{\mathrm{RFP}}$) and microscopy curve ($\chi$) are different, the relation $\chi_{\mathrm{RFP}} RFP = \chi N_{\mathrm{TF}}$ will be true according to the thermodynamic model of simple repression:

$$\text{fold-change}_{+1} = \frac{1}{1 + \chi_{\mathrm{RFP}}\mathrm{RFP}} = \frac{1}{1 + \chi N_{\mathrm{TF}}}. \tag{S1}$$

This leads to the following interpretation of the quantity $\chi_{\mathrm{RFP}} RFP$:

$$\chi_{\mathrm{RFP}} RFP = \lambda_{\mathrm{RFP}} \frac{1 + P\beta}{1 + P} RFP = \lambda_{\mathrm{mic}} \frac{1 + P\beta}{1 + P} \mu RFP = \chi N_{\mathrm{TF}}. \tag{S2}$$

Here the parameter $\lambda$ represents the binding affinity of CpxR with the subscript denoting the units it was measured in. The term $\mu$ is the scaling factor in units of TF per RFP signal and the parameters $\chi_{\mathrm{RFP}}$ and $\chi_{\mathrm{mic}}$ represent the rate of approach of in terms of their respective units. We find

$\chi_{\mathrm{RFP}} = 2.33 \times 10^{-4}$ (magenta curve) and $\chi = 1.3 \times 10^{-4}$ with the value of $\mu = 1.8$. We used this value of $\mu$ and multiplied the RFP signal in the cytometry data to scale the $x$-axis in Figure 5C and Figure 6E.

**Inference Plots for CpxR position sweep Data - Posterior distributions for the $\mathbf{FC_{max}}$ and $\boldsymbol{K}$**

As detailed in the Methods section, we used a particular formulation of the thermodynamic model [3] that used the TF abundance measurements from cytometry to rewrite the fold-change data for a given regulatory position as a function of the fold-change at the +1 position (see the Methods section on the Position Manifold Derivative). The benefit of this re-formulation was the ability to remove the binding affinity parameter and find the "intrinsic" regulatory parameters of the TF ($\alpha$ and $\beta$).

To extract $\alpha$ and $\beta$ for a given regulatory position, we began by inferring the $\mathrm{FC_{max}}$ and the $K = 1 + p\beta$ parameters using a Bayesian Sampling approach ([4]) as detailed in the inference procedure (see Methods Section for details). Figure S2 shows the result of the inference of the posterior distribution for the two parameters for each of the 12 regulatory positions. The first and third column shows the outcome of the Bayesian approach to sampling the posterior distribution for $\mathrm{FC_{max}}$ and $K$ parameters, and the second and fourth column presents the resulting transformation into the $\alpha$ and $\beta$ joint distribution space. The plots are arranged in order from highest $\beta$ at the top left of the figure to smallest $\beta$ at the bottom right. As seen, there is tight coupling for the joint distributions between the $\mathrm{FC_{max}}$ and the $K$ with an inverse dependence between the two parameters for activation (high values of $\mathrm{FC_{max}}$ are sampled jointly with low values of $K$ and vice versa) and opposite for the single repressing position $-54$ (low values of $\mathrm{FC_{max}}$ are sampled jointly with high values of $K$ and vice versa). Arranging them in order of decreasing $\beta$ serves to demonstrate an important point: the correlation between inference of $\alpha$ and $\beta$ becomes stronger as $\beta$ gets smaller (i.e. as $K$ approaches 1). In all cases, we can infer $\mathrm{FC_{max}}$ effectively but the individual value of one regulatory parameter depends strongly on the other because when $\beta P$ is small, $\mathrm{FC_{max}} \approx \alpha\beta$ and $K \approx 1$, and thus $\beta$ and $\alpha$ have a strong inverse relationship. When plotted on log axis this appears as a straight line (of slope $-1$) and the domain of $\beta$ from the inference sampling becomes less constrained. For all but one of the 12 regulatory positions ($-74$), we are unable to separate $\alpha$ and $\beta$ with any certainty and the inferred values cover a huge range that give the appropriate $\mathrm{FC_{max}}$.

**Testing alternative models of transcriptional regulation for the CpxR position sweep data**

In this section, we present an alternate interpretation of the CpxR position sweep data. Specifically we explore if the data can be explained by a simpler model with only one unique regulatory parameter for a given position manifold dataset. Specifically, the models we will evaluate in this section will assume the TF operates only through (de)stabilization of RNAP ($\alpha = 1$).

For the model inference runs, we set the energy of the promoter used in our synthetic circuit (DL5 promoter sequence [5]) as a global parameter across the 12 regulatory positions and allowed each position to infer its own stabilization ($\beta$) value, with $\mathrm{FC}_{\mathrm{max}}$ and the $K$ terms as defined in the position manifold section:

$$\mathrm{FC}^{y} = \frac{1+\mathrm{FC}_{\mathrm{max}}^{(y)} K\left(\frac{1-\mathrm{FC}^{(+1)}}{\mathrm{FC}^{(+1)}}\right)}{1+K\left(\frac{1-\mathrm{FC}^{(+1)}}{\mathrm{FC}^{(+1)}}\right)}, \tag{S3}$$

$$\mathrm{FC}_{\mathrm{max}}^{(y)} = \frac{\beta^{(y)}}{1+\left(\frac{\mathrm{P}}{1+\mathrm{P}}\right)\left(\beta^{(y)}-1\right)}, \tag{S4}$$

$$K = 1+\mathrm{P}\beta^{(y)}, \tag{S5}$$

where $\mathrm{FC}^{(y)}$ and $\mathrm{FC}_{\mathrm{max}}^{(y)}$ represent the fold-change and $\mathrm{FC}_{\mathrm{max}}$ effective parameter when the binding site is introduced at position $y$ on the promoter. Note that $\mathrm{FC}_{\mathrm{max}}$ no longer has the acceleration parameter. For the Bayesian inference scheme, we set the prior of the DL5 promoter sequence energy as a uniform distribution between the values $-10\ k_{\mathrm{B}}T$ and $-2\ k_{\mathrm{B}}T$. The sequence energetics have been measured in prior work as $-6.5\ k_{\mathrm{B}}T$, as such we believed this was a reasonable interval for the chain to sample. For the position dependent stabilization parameters, we used a uniform distribution with acceptable bounds for all activation positions. We modeled the likelihood function as product of normal distributions (assuming the global likelihood function is a product of the individual position specific likelihood functions) with the mean of these position specific distributions as the theoretical fold-change value generated from the thermodynamic model conditioned on the parameters. To ensure that differences in the fold-change profiles between strong activation positions and weak activation positions was adequately conveyed in the likelihood function, we made the standard deviation for the respective position specific normal distributions a hyperparameter in our model, with the final form of the likelihood function as follows:

$$\prod_{i}^{\text{Position}} \prod_{j}^{\text{Data Points}} \text{Normal}\left(\text{FC}^{(i)}_{\text{stabilization}}\left(\text{P}, \beta^{i}\right) | u = FC^{i}_{j}, sd = \sigma^{(i)})\right), \quad \text{(S6)}$$

and:

$$\text{FC}^{(i)}_{\text{stabilization}} = \frac{1 + \text{FC}^{(i)}_{\max}\left(\text{P}, \beta^{(i)}\right) K\left(\text{P}, \beta^{(i)}\right) \left(\frac{1-\text{FC}^{(+1)}}{\text{FC}^{(+1)}}\right)}{1 + K\left(\text{P}, \beta^{(i)}\right) \left(\frac{1-\text{FC}^{(+1)}}{\text{FC}^{(+1)}}\right)}. \quad \text{(S7)}$$

We initialized the inference procedure as sampling from a global vector, $\theta$, that contained the global and position specific parameters in our model,

$$\theta = \left[\ \text{P}^{\text{energy}}, \beta^{\text{-48}}, \beta^{\text{-50}}, \beta^{\text{-54}} ..., \beta^{\text{-80}}, \beta^{\text{-82}}.\ \right] \quad \text{(S8)}$$

The inference procedure was run for 50000 runs initialized on 4 different chains to ensure adequate sampling of the joint parameter space. The fits for the "stabilization only" model are shown in Figure S3. In this figure, we plot the results of the thermodynamic model from the inference sampling. It is clear that the stabilization only model fails in capturing the highest activation position ($-64$) and the curvature seen in some positions ($-50$, $-54$). This failure to explain strong activation is expected, as in a model without acceleration ($\alpha = 1$) the maximum possible fold-change is constrained by the individual occupancy of the promoter by RNAP (the maximal possible fold-change is roughly $1/P$ for weak promoters or more precisely $(1+P)/P$ if the weak promoter assumption is lifted [6]); intuitively, in this model without acceleration if the constitutive promoter has polymerase occupancy 10% of the time, the largest fold-change possible is 10 (corresponding to 100% occupancy). To reach the fold-change values obtained in the CpxR position sweep data, the inferred value of $P$ needs to be at least $\sim 10$ fold lower than the value of $P$ measured in previous work [5]. As such, we feel confident that this model can not sufficiently describe our data.

It is worth noting that even if this assumption is incorrect and in reality $P$ is much smaller than we expect from previous measurements, the model without acceleration still does not describe the data well; the theory does not match the curvature of the data seen in some positions such as $-50$, $-54$, and $-60$ (Figure S3). This feature highlights the importance of acceleration ($\alpha$) in explaining these positions and the regulation data at large.

**Physiological effects of TF titration on synthetic circuit expression**

One concern for the gene expression measurements was separating the fundamental regulatory role of a TF from the apparent expression changes due to potential physiological effects such as slowing growth rates from, for instance, high inducer concentrations or changing TF concentration in the cell [7, 8, 9] brought about by inducing the TFs to different levels. Increasing concentrations of the TF in the cell could potentially alter YFP expression by turning on or off genes involved in global regulation of translation or through a host of post-transcriptional events. In Figure S4, we do not see a major perturbation in synthetic circuit expression for most of the TF titration strains as seen in Figure S4 where YFP expression hovers at the FC=1 line as the TF concentration increases.

**Robustness of the Position Manifold Results**

To analyze the fold-change data of our experiments we bin the single-cell fluorescence measurements to find the average fold-change of cells with similar TF concentrations (mCherry levels). To accomplish this we divided the data into a specified number of bins based on the proportion of the total data points and calculated the ensemble fold-change from the cells in each bin. In Figure S5 and Figure S6 we show how the determination of the parameters $\beta$ and $\alpha$ depends on this choice. In these plots the inferred value of alpha and beta for each of the 12 positions are shown for 14 different bin numbers (6, 8, 10, 12, 14, 16, 18, 20, 24, 26, 28, 30, 32, 36) number of bins and plotted against the value found with 22 bins (used in the main text). The quatity on the y-axis is a measurement of the degree to which the values of $\beta$ (or $\alpha$) differ from the reference bin across all regulatory positions, and is computed by taking the mean of the $\log_{10}$ ratios between the reference bin and the bin under consideration for each of the 12 regulatory position. In Figure S5, we see that the inferred value of $\beta$ for other bin sizes is tight across most of the regulatory positions with more substantial deviations as the number of bins becomes very small ($< 12$). This phenomenon is an indication to the degree the larger bin sizes (smaller bin numbers) inadequately convey the curvature inherent in the data, pushing more of the regulatory positions to overstimate the degree of curvature for certain regulatory positions. Figure S6 shows this same measure for the inferred values of $\alpha$, where we we once again see consistency in the inferred value across most of the regulatory positions except for the small number of bins as seen in Figure S5. Crucially, the inference of $\alpha$ and $\beta$ is not critically sensitive to the choice of bin size above 12 bins.

**List of Strains and primers used for the TF-titration cloning and measurements**

**Primers for amplifying the TF gene cassettes from the MG1655 genome**

The following primers (Table S1) were used to amplify the coding sequence (without the stop codon) of the 6 TFs profiled from the MG1655 genome. The amplified coding regions were then cloned by Gibson assembly into the pTet-TF-AEK-mcherry that has the $P_{Tet}$ promoter in frame with the flexible AEK linker sequence (GCAGAAGCAGCAGCAAAGGAAGCAGCAGCAAAG-GCA) and mCherry sequence. The overhangs for the insertion into the pTet-TF-AEK-mCherry plasmid were included at the 5' end of the primer sequence followed by the remainder sequence that served as the priming sequence in the MG1655 genome beginning with the start codon for the TF sequence. The resulting plasmid served as a template for chromosomal integration of the TF-AEK-mCherry fusion at the *ybcN* locus. Primers for chromosomal integration at the *ybcN* locus is described in [1].

**TF strains harboring the inducible gene circuit for control of TF copy number in *E. coli***

The following table lists the engineered strains that allowed for systematic titration of TF copy number for the 6 TFs presented in this work. Details on how the TF-mCherry fusion strains were generated are found in the Methods section under the heading "Engineering the titratable TF-mCherry fusion strain". Each TF-fusion strain was made in the MG1655 *E. coli* genotype, with the integration of the constitutively expressed TetR cassette integrated in the *gspI* locus (Table S2). The constitutive strain was engineered by P1-transduction of the KEIO strain harboring the deletion of the TF (KO strain) into the MG1655 genotype with the *gspI* locus integration.

**TF Binding Sites used for assessing the +1 and -61 regulatory profile**

The following binding sites listed in Table 2 were cloned into the synthetic gene circuit integrated at the *galK* locus. The synthetic circuit contained a defined promoter sequence with a previously characterized variant of the *LacUV5* core promoter sequence [5] and amplified for integration into the *galK* locus. The cloning of these TF binding sequences (Table S3) at the defined upstream and downstream locations at the promoter was done as specified in the Methods Section with the strains listed in Table S4.

**Position Sweep Primers used for CpxR Position Sweep Cloning**

The capitalized nucleotides represent the priming sequence for the position-specific amplification of the $P_{DL5}$ upstream promoter sequence. Primers were designed using a custom Python script that found an optimal sequence to amplify the promoter sequence at a defined location relative to the +1 with both forward and reverse primers designed to have comparable melting temperatures (Table S5). The initial sequence in the 5' region represents overhangs for the pDONR P4-P1r plasmid that contained the *ccdb* toxin cassette (the sequence is the reverse complement of the primers amplifying the region and the resulting amplicons from both plasmids were used to clone the $P_{DL5}$ position specific strains using Gibson Assembly. All positions are designated based on the $3'$ end of the TF binding sequence.

**Positional Cloning Strains**

The following strains were generated from the cloning procedure described in the Methods section "Cloning the TF specific binding location gene circuits". The primers from Table S5 were used to generate these strains with the positional notation determined by the location of the 3' end of the TF binding sequence after the digestion and ligation cloning as described in the Methods Section. These plasmids are listed in Table S6.

**List of TF strains used in the Microscopy Experiments (Survey of the regulatory activity of 6 TFs at +1 and -61)**

To profile the regulatory response of the 6 TFs at positions +1 and -61 relative to the transcription start site (TSS), we used the Engineered TF fusion strains listed in Table S2 and substituted the *galK* locus with the synthetic circuit derived from amplifying the regulatory sequence harboring the promoter, YFP coding region, and transcriptional terminator elements from the $P_{DL5}$ plasmid (Table S4). The primers amplifying the regulatory sequence had overhangs for the *galK* locus and the integration protocol is described in the Methods Section. These strains are listed in Table S7.

**List of CpxR position sweep strains)**

The following CpxR position regulation strains were made by using the positional cloning plasmids listed in Table S6 according to the digestion and ligation procedure detailed in the Methods Section "Cloning the TF specific binding location gene circuits". These plasmids were then transformed into the relevant geneotypes (CpxR KO and CpxR-mCherry) to measure the regulatory profile of the TF activity. These strains are listed in Table S8.

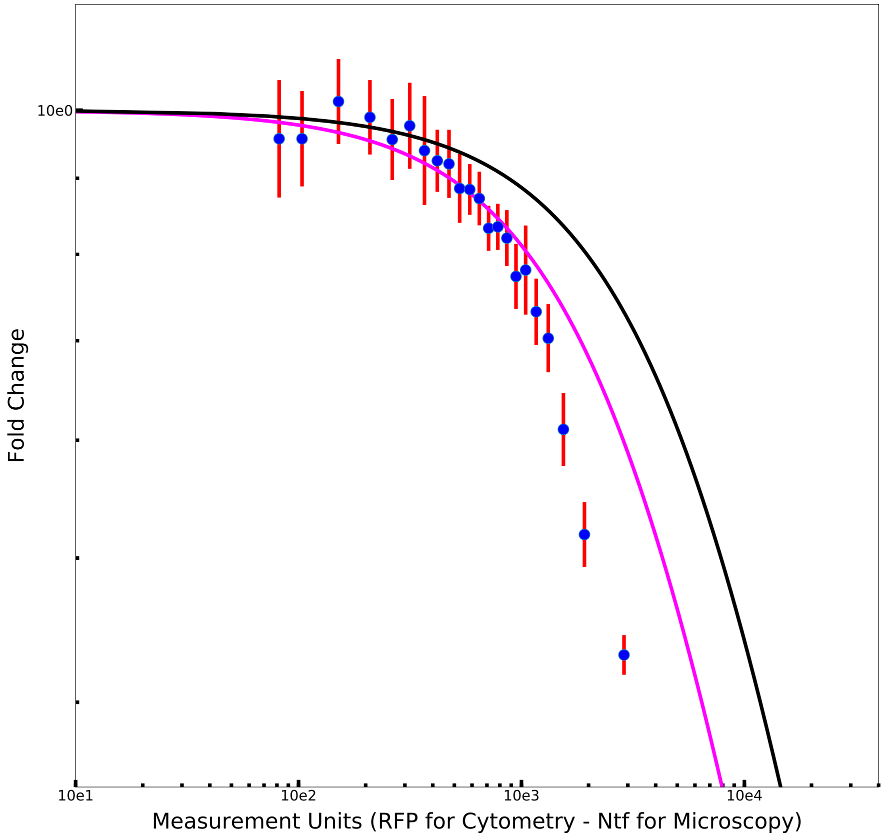


Figure S1: **Plot of the fold-change in regulation for the +1 binding site from the cytometry and microscopy approach.** We used the +1 data measured on microscopy and cytometry to generate the scaling factor that would allow for conversion of arbitrary fluoresence in cytometry to TF copy number. The data points represent the measured fold-change values from the cytometry approach and the magenta line represents the fit generated from the thermodynamic model where $\chi_{\text{RFP}}$ is fitted to the global CpxR position regulation dataset (See Methods Section "Using the position manifold parameters to generate the thermodynamic model in FC vs $N_{\text{TF}}$ space" for details). The black line represents the model where $\chi$ is fit to the microscopy data. Note that the $x$-axis for the cytometry curve and data points is in RFP units while for the microscopy curve it is in TF copy number. The difference between the curves can be removed by scaling the value of $\chi_{\text{RFP}}$ by $\chi$, and this resulting scaling value ($\mu = 1.8$ TFs/RFP unit) is the factor that converts the RFP measurements to TF copy number.

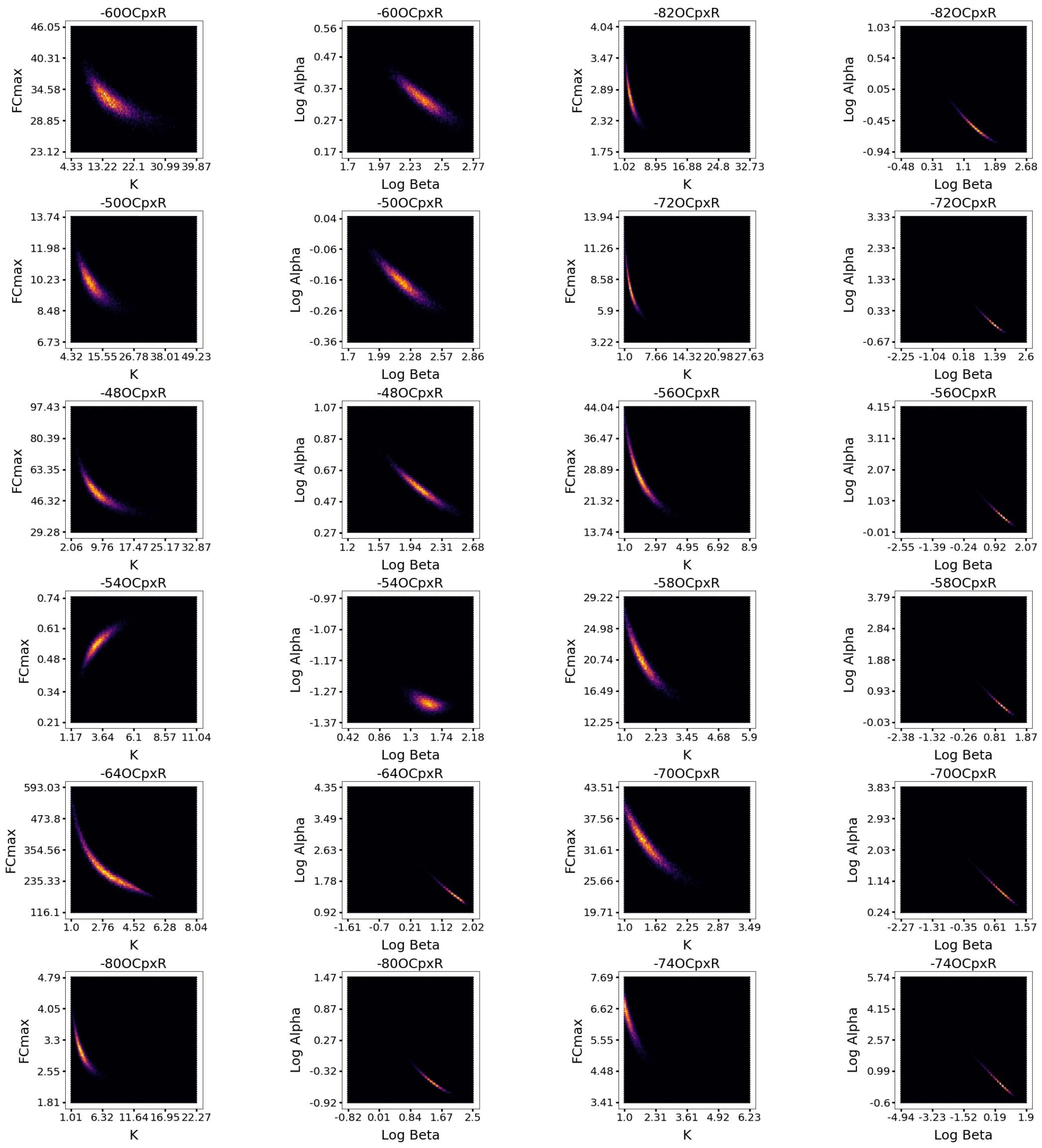


Figure S2: **Inference plots for the Stabilization Only Model.** Plots of the $\mathrm{FC}_{\max}$ and $K$ from the model inference of the position manifold approach in the first column with the second columns representing the transformation into the joint space of $\alpha$ and $\beta$ as presented in the main text. The positions are arranged from highest activation to moderate repression. Note the strong inverse dependence between the $\mathrm{FC}_{\max}$ and $K$ parameters for the activation positions that is reversed for the repression position $-54$. A log-log plot of the $\alpha$ and $\beta$ parameters will yield a straight line if the $K$ is 1, and the sampled space for $\alpha$ and $\beta$ will be large preventing inference on the mode of regulation for the given regulatory position.

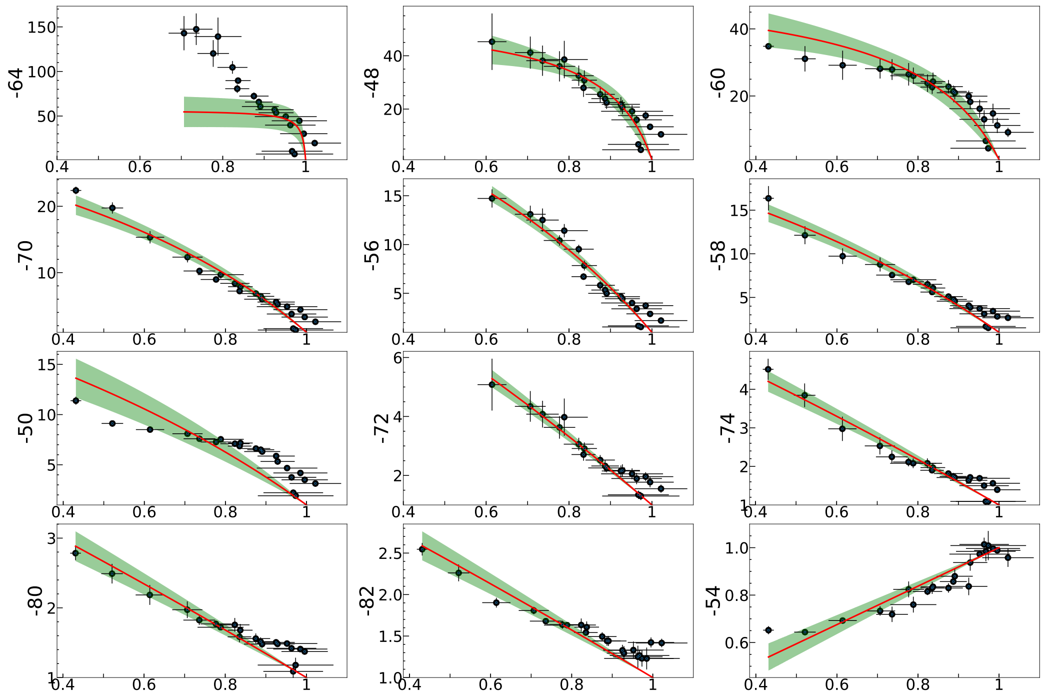


Figure S3: **Plots for the Stabilization Only Model.** The position plots are arranged from highest to lowest activation positions. The regulatory contours are generated from the inferred parameters according to a model that lacks acceleration ($\alpha = 1$). The black data points are from the position manifold approach as described in the Main Text. The red solid line represents the model expectation conditioned on the inferred parameters (the position specific $\beta$ and the global $P$ parameter) with the green shaded area as 2 standard deviations from this expectation. Note that the stabilization model fails to capture the data for positions with strong curvature as well as the highest activation position ($-64$).

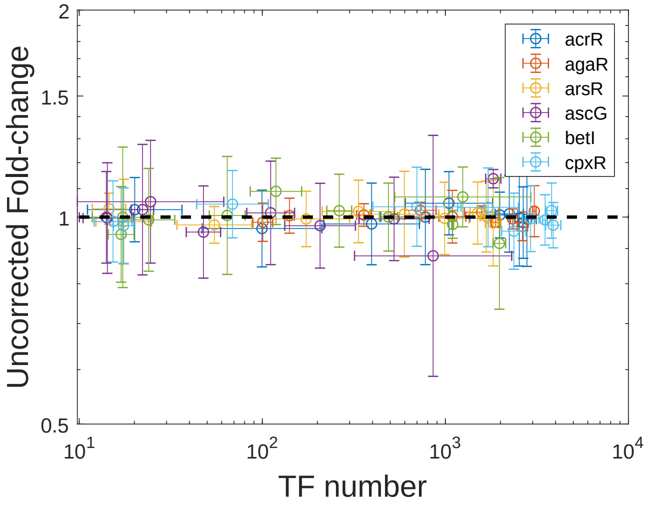


Figure S4: **Physiology effects of TF-titration on YFP expression.** TF-titration strains regulating the control synthetic circuit (*Psz2-DelBs*) as measured using microscopy. The dashed black line represents FC=1, and we see most of the TFs do not appreciably alter YFP expression simply through the titration effect.

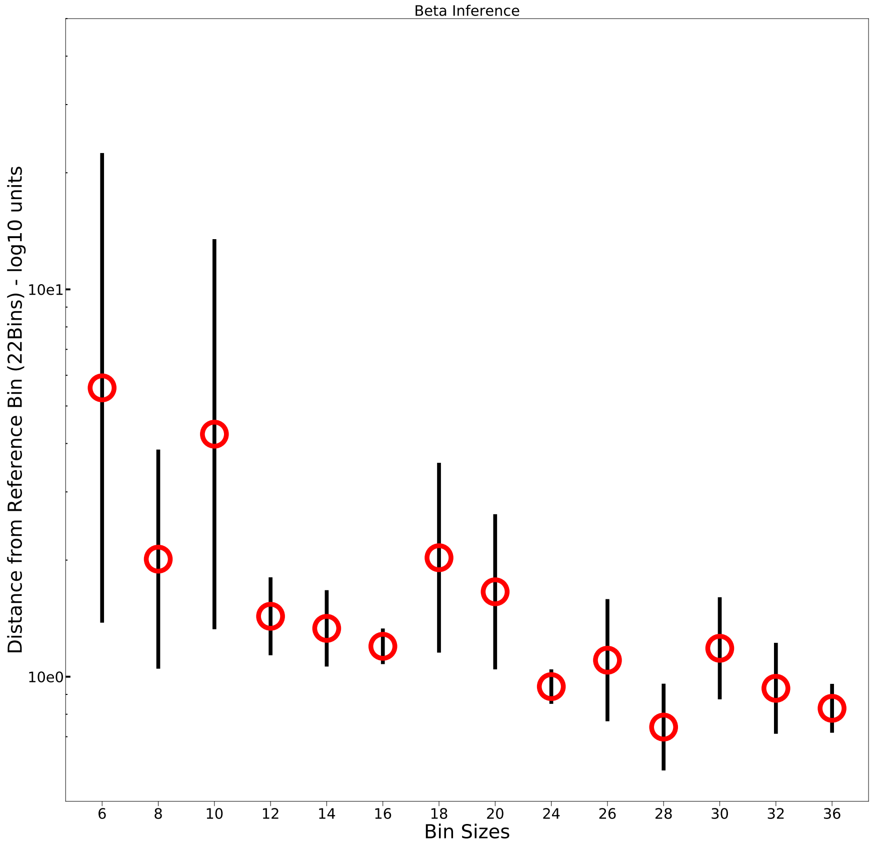


Figure S5: **Deviations in the median of the stabilization parameter $\beta$ as the number of bins changes.** The position manifold approach was used on the 12 CpxR regulatory positions with different number of proportional bins and the median value of $\beta$ for these 12 positions was compared to the bin number used in the main text. The $y$-axis represents the distance in the median parameter values relative to this refernece bin for bin numbers that span from 6 to 36 as detailed in the corresponding SI section. Note that for smaller bins ($< 12$) the deviations are noticeable as the larger bin size masks the appropriate degree of curvature in the datasets, leading to an overestimation of $\beta$ in these cases.

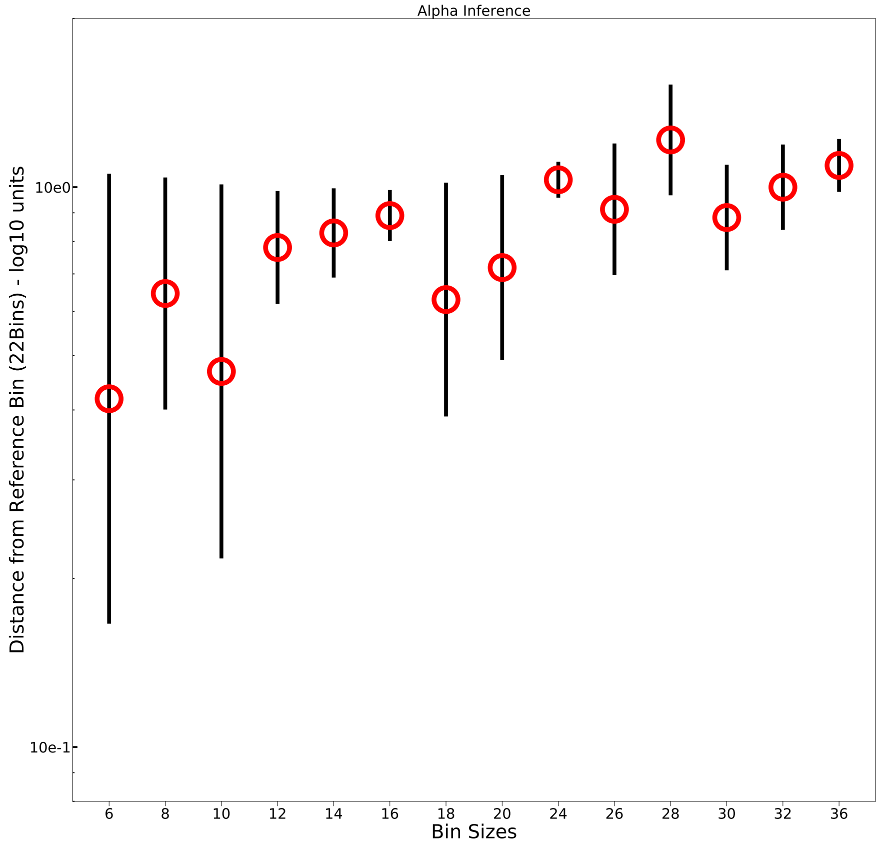


Figure S6: **Robustness of the acceleration ($\alpha$) values as the bin size is changed for position manifold binning.** Corresponding plot to SI Figure 5 for the difference in the median $\alpha$ values for different bin sizes compared to the reference bin. Note that as the values of $\beta$ in the lower bin numbers ($< 12$) were overestimated, the lower median values of $\alpha$ for these bin sizes are therefore expected. The inferred median value of $\alpha$, however, is fairly robust at larger bin numbers ($> 12$)

| TF | Primer Sequence |
|---|---|
| AcrR FP | CAT TAA AGA GGA GAA AGG TAC CAT ATG GCA CGA AAA ACC AAA CAA GAA GCG |
| AcrR RP | GCT GCT GCT TCC TTT GCT GCT GCT TCT GCT TCG TTA GTG GCA GGA TTA CGA AGC G |
| AgaR FP | CAT TAA AGA GGA GAA AGG TAC CAT ATG AGT AAT ACC GAC GCT TCA GG |
| AgaR RP | GCT TCC TTT GCT GCT GCT TCT GCC TCC CCG ACC AGA ATC ACT TCA ACC |
| ArsR FP | CAT TAA AGA GGA GAA AGG TAC CAT ATG TCA TTT CTG TTA CCC |
| ArsR RP | GCT GCT GCT TCC TTT GCT GCT GCT TCT GCA CTG CAA ATG TTC TTA CTG TCC C |
| AscG FP | CAT TAA AGA GGA GAA AGG TAC CAT ATG ATG ACG ACG ATG CTG GAA GTG G |
| AscG RP | GCT TCC TTT GCT GCT GCT TCT GCT CGC GAA GGA GCA ATG AGT G |
| BetI FP | CAT TAA AGA GGA GAA AGG TAC CAT ATG CCC AAA TTG GGG ATG C |
| BetI RP | GCT TCC TTT GCT GCT GCT TCT GCA TCG GTG GGT AGA TGC TGA G |
| CpxR FP | CAT TAA AGA GGA GAA AGG TAC CAT ATG AAT AAA ATC CTG TTA GTT G |
| CpxR RP | GCT TCC TTT GCT GCT GCT TCT GCT GAA GCA GAA ACC ATC AGA TAG C |

Table S1: **Primers for amplifying the TF gene cassettes from the MG1655 genome** FP refers to the forward primer sequence and RP refers to the reverse primer sequence. The primer pairs were designed to amplify the TF gene without the translational stop codon to allow for the TF-fusion to the mCherry gene through a linker sequence.

| **Engineered TF fusion Strains** | **Genotype** |
|---|---|
| AcrR (Constitutive) | MG1655, AcrRKO, *gspI*<>1pn25-TetR |
| AcrR (Regulation) | *gspI*<>1pn25-TetR, *ybcN*<>3*1-AcrR-AEK-mCheery, AcrRKO |
| AgaR (Constitutive) | AgaRKO, *gspI*<>1pn25-TetR |
| AgaR (Regulation) | *gspI*<>1pn25-TetR, *ybcN*<>3*1-AgaR-AEK-mCheery, AgaRPKO |
| ArsR (Constitutive) | ArsRKO, *gspI*<>1pn25-TetR |
| ArsR (Regulation) | *gspI*<>1pn25-TetR, *ybcN*<>3*1-ArsR-AEK-mCheery, ArsRKO |
| AscG (Constitutive) | AscGKO, *gspI*<>1pn25-TetR |
| AscG (Regulation) | *gspI*<>1pn25-TetR, *ybcN*<>3*1-AscG-AEK-mCheery, AscGKO |
| BetI (Constitutive) | BetIKO, *gspI*<>1pn25-TetR |
| BetI (Regulation) | *gspI*<>1pn25-TetR, *ybcN*<>3*1-BetI-AEK-mCheery, BetIKO |
| CpxR (Constitutive) | C pxRKO, *gspI*<>1pn25-TetR |
| CpxR (Regulation) | *gspI*<>1pn25-TetR, *ybcN*<>3*1-cpxR-AEK-mCheery, cpxRKO |

Table S2: **Titration strains to control the average copy number of the 6 TFs studied here.** All strains had the TF-mCherry expression cassette integrated into the *ybcN* locus.

| TF | Promoter containing sequence | Binding Sequence |
|---|---|---|
| AcrR | $P_{acrR}$ | TACATACATTCACAAATGTATGTA |
| AgaR | $P_{kbaZ}$ | CTTTCGTTTCATTTCGTTT |
| ArsR | $P_{arsR}$ | TAAGTCATATATGTTTTTGACTTA |
| AscG | $P_{ascF}$ | TGAAACCGGTTTCT |
| BetI | $P_{betI}$ | TATATTGAACGTCCAATCAA |
| CpxR | $P_{ppiA}$ | GTAAAATTAGGTAAA |

Table S3: **Binding sequences for the 6 TFs adapted from the endogenous regulated promoters in *E. coli*.**

| +1 **Binding Position** | −61 **Binding Position** | **Parent Strain** |
|---|---|---|
| *galK*<>25PDL5AcrRbs+1-YFP | *galK*<>25PDL5AcrRbs-61-YFP | *gspI*<>1pn25-TetR; MG1655 |
| *galK*<>25PDL5 AgaRbs +1-YFP | *galK*<>25PDL5 AgaRbs-61-YFP | *gspI*<>1pn25-TetR; MG1655 |
| *galK*<>25PDL5 ArsRbs +1-YFP | *galK*<>25PDL5 ArsRbs-61-YFP | *gspI*<>1pn25-TetR; MG1655 |
| *galK*<>25PDL5 AscGbs+1-YFP | *galK*<>25PDL5 AscGbs-61-YFP | *gspI*<>1pn25-TetR; MG1655 |
| *galK*<>25PDL5 BetIbs+1-YFP | *galK*<>25PDL5 BetIbs-61-YFP | *gspI*<>1pn25-TetR; MG1655 |
| *galK*<>25PDL5 CpxRbs +1-YFP | *galK*<>25PDL5 CpxRbs-61-YFP | *gspI*<>1pn25-TetR; MG1655 |

Table S4: **Strains for measuring regulation at** +1 **and** −61 **for 6 TFs.** Synthetic circuits designed by cloning the binding sites for the 6 TFs listed in Table S3 at 2 distinct positions in the $P_{DL5}$ promoter sequence.

| **Position Relative to TSS** | **Forward Primer** | **Reverse Primer** |
|---|---|---|
| −41 | ctccgaagacctcgtgTTTACCCTTTATGCTTCCGGCT | gaattcgaagactggctcCACGAGGTGAAGCACGA |
| −43 | ctccgaagacctcgtgTGTTTACCCTTTATGCTTCCGG | gaattcgaagactggctcCGAGGTGAAGCACGAAAGG |
| −45 | ctccgaagacctcgtgCGTGTTTACCCTTTATGCTTCCG | gaattcgaagactggctcAGGTGAAGCACGAAAGGG |
| −47 | ctccgaagacctcgtgCTCGTGTTTACCCTTTATGCTTCC | gaattcgaagactggctcGTGAAGCACGAAAGGGC |
| −49 | ctccgaagacctcgtgACCTCGTGTTTACCCTTTATGC | gaattcgaagactggctcGAAGCACGAAAGGGCCTC |
| −51 | ctccgaagacctcgtgTCACCTCGTGTTTACCCTTTATGC | gaattcgaagactggctcAGCACGAAAGGGCCTCGT |
| −53 | ctccgaagacctcgtgCTTCACCTCGTGTTTACCCTTTATG | gaattcgaagactggctcCACGAAAGGGCCTCGTG |
| −55 | ctccgaagacctcgtgTGCTTCACCTCGTGTTTACC | gaattcgaagactggctcCGAAAGGGCCTCGTGATAC |
| −57 | ctccgaagacctcgtgCGTGCTTCACCTCGTGTTTAC | gaattcgaagactggctcAAAGGGCCTCGTGATACG |
| −59 | ctccgaagacctcgtgTTCGTGCTTCACCTCGTG | gaattcgaagactggctcAGGGCCTCGTGATACGC |
| −61 | ctccgaagacctcgtgCTTTCGTGCTTCACCTCGT | gaattcgaagactggctcGGCCTCGTGATACGCCTAT |
| −63 | ctccgaagacctcgtgCCCTTTCGTGCTTCACC | gaattcgaagactggctcCCTCGTGATACGCCTATTTCG |
| −65 | ctccgaagacctcgtgGGCCCTTTCGTGCTTCAC | gaattcgaagactggctcTCGTGATACGCCTATTTCGATG |
| −67 | ctccgaagacctcgtgGAGGCCCTTTCGTGCTTC | gaattcgaagactggctcGTGATACGCCTATTTCGATGGG |
| −69 | ctccgaagacctcgtgACGAGGCCCTTTCGTGC | gaattcgaagactggctcGATACGCCTATTTCGATGGGTTAATG |
| −73 | ctccgaagacctcgtgTATCACGAGGCCCTTTCGTG | gaattcgaagactggctcCGCCTATTTCGATGGGTTAATGT |
| −75 | ctccgaagacctcgtgCGTATCACGAGGCCCTTTC | gaattcgaagactggctcCCTATTTCGATGGGTTAATGTCATGG |
| −81 | ctccgaagacctcgtgAATAGGCGTATCACGAGGCCCTT | gaattcgaagactggctcTCGATGGGTTAATGTCATGGAG |
| −87 | ctccgaagacctcgtgCATCGAAATAGGCGTATCACGAG | gaattcgaagactggctcGGTTAATGTCATGGAGCTAATGGT |
| −93 | ctccgaagacctcgtgTTAACCCATCGAAATAGGCGTATC | gaattcgaagactggctcTGTCATGGAGCTAATGGTTTCTTAG |
| −99 | ctccgaagacctcgtgATGACATTAACCCATCGAAATAGGC | gaattcgaagactggctcGGAGCTAATGGTTTCTTAGACGTCG |
| −105 | ctccgaagacctcgtgAGCTCCATGACATTAACCCATC | gaattcgaagactggctcAATGGTTTCTTAGACGTCGGATG |

Table S5: **Primers used in cloning the *ccdB* cassette at the upstream and downstream locations of the $P_{DL5}$ promoter sequence.**

| ***ccdB* Cloning Strains** |
| --- |
| pZS25LongUPDL5-41-ccdB-YFP |
| pZS25LongUPDL5-53-ccdB-YFP |
| pZS25LongUPDL5-55-ccdB-YFP |
| pZS25LongUPDL5-57-ccdB-YFP |
| pZS25LongUPDL5-67-ccdB-YFP |
| pZS25LongUPDL5-69-ccdB-YFP |
| pZS25LongUPDL5-73-ccdB-YFP |
| pZS25LongUPDL5-43-ccdB-YFP |
| pZS25LongUPDL5-47-ccdB-YFP |
| pZS25LongUPDL5-49-ccdB-YFP |
| pZS25LongUPDL5-51-ccdB-YFP |
| pZS25LongUPDL5-65-ccdB-YFP |
| pZS25LongUPDL5-105-ccdB-YFP |
| pZS25LongUPDL5-87-ccdB-YFP |
| pZS25LongUPDL5-93-ccdB-YFP |
| pZS25LongUPDL5-59-ccdB-YFP |
| pZS25LongUPDL5-63-ccdB-YFP |
| pZS25LongUPDL5-75-ccdB-YFP |
| pZS25LongUPDL5-81-ccdB-YFP |
| pZS25LongUPDL5-99-ccdB-YFP |
| pZS25LongUPDL5-61-ccdB-YFP |

Table S6: ***ccdB* plasmids for rapid cloning.** Plasmids used to clone the *ppiA* binding sequence for the assessing the position dependent regulatory profiles of CpxR.

| **TF-Fusion Strain** | $+1$ − **Synthetic Circuit Strains** | $-61$ − **Synthetic Circuit Strains** | **No Binding Sequence − Synthetic Circuit Strains** |
|---|---|---|---|
| AcrR (Constitutive) | AcrRKO, *gspI*<>1pn25-TetR, *galK*<>25DL5AcrRbs+1-YFP | AcrRKO, *gspI*<>1pn25-TetR, *galK*<>25DL5AcrRbs-61-YFP | AcrRKO, *gspI*<>1pn25-TetR, *galK*<>25DL5-YFP |
| AcrR (Regulation) | *gspI*<>1pn25-TetR, *ybcN*<>3*1-AcrR-AEK-mCheery, *galK*<>25DL5AcrRbs+1-YFP, AcrRKO | *gspI*<>1pn25-TetR, *ybcN*<>3*1-AcrR-AEK-mCheery, *galK*<>25DL5AcrRbs-61-YFP, AcrRKO | *gspI*<>1pn25-TetR, *ybcN*<>3*1-AcrR-AEK-mCheery, *galK*<>25DL5-YFP, AcrRKO |
| AscG (Constitutive) | AscGKO, *gspI*<>1pn25-TetR, *galK*<>25DL5AscGbs+1-YFP | AscGKO, *gspI*<>1pn25-TetR, *galK*<>25DL5AscGbs-61-YFP | AscGKO, *gspI*<>1pn25-TetR, *galK*<>25DL5-YFP |
| AscG (Regulation) | *gspI*<>1pn25-TetR, *ybcN*<>3*1-AscG-AEK-mCheery, *galK*<>25DL5AscGbs+1-YFP, AscGKO | *gspI*<>1pn25-TetR, *ybcN*<>3*1-AscG-AEK-mCheery, *galK*<>25DL5AscGbs-61-YFP, AscGKOs | *gspI*<>1pn25-TetR, *ybcN*<>3*1-AscG-AEK-mCheery, *galK*<>25DL5-YFP , AscGKO |
| AgaR (Constitutive) | AgaRKO, *gspI*<>1pn25-TetR, *galK*<>25DL5AgaRbs+1-YFP | AgaRKO, *gspI*<>1pn25-TetR, *galK*<>25DL5AgaRbs-61-YFP | AgaRKO, *gspI*<>1pn25-TetR,*galK*<>25DL5-YFP |
| AgaR (Regulation) | *gspI*<>1pn25-TetR, *ybcN*<>3*1-AgaR-AEK-mCheery, *galK*<>25DL5AgaRbs+1-YFP, AgaRKO | *gspI*<>1pn25-TetR, *ybcN*<>3*1-AgaR-AEK-mCheery, *galK*<>25DL5AgaRbs-61-YFP, AgaRKO | *gspI*<>1pn25-TetR, *ybcN*<>3*1-AgaR-AEK-mCheery, *galK*<>25DL5-YFP, AgaRKO |
| ArsR (Constitutive) | ArsRKO, *gspI*<>1pn25-TetR, *galK*<>25DL5ArsRbs+1-YFP | ArsRKO, *gspI*<>1pn25-TetR,*galK*<>25DL5ArsRbs-61-YFP | ArsRKO, *gspI*<>1pn25-TetR,*galK*<>25DL5-YFP |
| ArsR (Regulation) | *gspI*<>1pn25-TetR, *ybcN*<>3*1-ArsR-AEK-mCheery, *galK*<>25DL5ArsRbs+1-YFP, ArsRKO | *gspI*<>1pn25-TetR, *ybcN*<>3*1-ArsR-AEK-mCheery, *galK*<>25DL5ArsRbs-61-YFP, ArsRKO | *gspI*<>1pn25-TetR, *ybcN*<>3*1-ArsR-AEK-mCheery, *galK*<>25DL5-YFP, ArsRKO |
| BetI (Constitutive) | BetIKO, *gspI*<>1pn25-TetR, *galK*<>25DL5BetIbs+1-YFP | BetIKO, *gspI*<>1pn25-TetR, *galK*<>25DL5BetIbs-61-YFP | BetIKO, *gspI*<>1pn25-TetR,*galK*<>25DL5-YFP |
| BetI (Regulation) | *gspI*<>1pn25-TetR, *ybcN*<>3*1-BetI-AEK-mCheery, *galK*<>25DL5BetIbs+1-YFP,BetIKO | *gspI*<>1pn25-TetR, *ybcN*<>3*1-BetI-AEK-mCheery, *galK*<>25DL5BetIbs-61-YFP,BetIKO | *gspI*<>1pn25-TetR, *ybcN*<>3*1-BetI-AEK-mCheery, *galK*<>25DL5-YFP BetIKO |
| CpxR (Constitutive) | CpxRKO, *gspI*<>1pn25-TetR, *galK*<>25DL5CpxRbs+1-YFP | CpxRKO, *gspI*<>1pn25-TetR,*galK*<>25DL5CpxRbs-61-YFP | CpxRKO, *gspI*<>1pn25-TetR, *galK*<>25DL5-YFP |
| CpxR (Regulation) | *gspI*<>1pn25-TetR, *ybcN*<>3*1-CpxR-AEK-mCheery, *galK*<>25DL5CpxRbs+1-YFP, CpxRKO | *gspI*<>1pn25-TetR, *ybcN*<>3*1-CpxR-AEK-mCheery, *galK*<>25DL5CpxRbs-61-YFP, CpxRKO | *gspI*<>1pn25-TetR, *ybcN*<>3*1-CpxR-AEK-mCheery, *galK*<>25DL5-YFP, CpxRKO |

Table S7: **Strains used to survey the regulatory activity of 6 TFs at $+1$ and $-61$.** All synthetic gene circuits were integrated in the *E. coli* genome, and the constitutive strains differ by the omission of the titratable TF-mCherry circuit integrated at the *ybcN* locus

| CpxR Position Sweep Plasmids |
|---|
| pZS25LongUPDL5+1OCpxR-YFP |
| pZS25LongUPDL5-41OCpxR-YFP |
| pZS25LongUPDL5-43OCpxR-YFP |
| pZS25LongUPDL5-47OCpxR-YFP |
| pZS25LongUPDL5-49OCpxR-YFP |
| pZS25LongUPDL5-51OCpxR-YFP |
| pZS25LongUPDL5-53OCpxR-YFP |
| pZS25LongUPDL5-55OCpxR-YFP |
| pZS25LongUPDL5-57OCpxR-YFP |
| pZS25LongUPDL5-59OCpxR-YFP |
| pZS25LongUPDL5-61OCpxR-YFP |
| pZS25LongUPDL5-63OCpxR-YFP |
| pZS25LongUPDL5-65OCpxR-YFP |
| pZS25LongUPDL5-67OCpxR-YFP |
| pZS25LongUPDL5-69OCpxR-YFP |
| pZS25LongUPDL5-73OCpxR-YFP |
| pZS25LongUPDL5-75OCpxR-YFP |
| pZS25LongUPDL5-81OCpxR-YFP |
| pZS25LongUPDL5-87OCpxR-YFP |
| pZS25LongUPDL5-93OCpxR-YFP |
| pZS25LongUPDL5-delBS -YFP |
| pZS25LongUPDL5-99OCpxR-YFP |
| pZS25LongUPDL5-105OCpxR-YFP |

Table S8: **Cpxr position sweep plasmids.** Plasmids were generated using the *ccdB* cloning vectors in Table S6. These position cloning vectors had the *ppiA* binding sequence swapped at the specified locations on the plasmid.
